# CtIP promotes the motor activity of DNA2 to accelerate long-range DNA end resection

**DOI:** 10.1101/789230

**Authors:** Ilaria Ceppi, Sean M. Howard, Kristina Kasaciunaite, Cosimo Pinto, Roopesh Anand, Ralf Seidel, Petr Cejka

**Affiliations:** Institute for Research in Biomedicine, Università della Svizzera italiana (USI), Faculty of Biomedical Sciences, Bellinzona, Switzerland; Department of Biology, Institute of Biochemistry, Eidgenössische Technische Hochschule (ETH), Zürich, Switzerland; Peter Debye Institute for Soft Matter Physics, Universität Leipzig, Leipzig, Germany; Institute of Molecular Cancer Research, University of Zürich, Zürich, Switzerland

**Author notes:** Lead Contact: Petr Cejka, Institute for Research in Biomedicine, Via Vincenzo Vela 6, 6500 Bellinzona, Switzerland.

## Abstract

BLM or WRN helicases function with the DNA2 helicase-nuclease to resect DNA doublestrand breaks and initiate homologous recombination. Upon DNA unwinding by BLM/WRN, RPA directs the DNA2 nuclease to degrade the 5’-strand, revealing the 3’ overhang needed for recombination. RPA bound to ssDNA also represents a barrier, explaining the need for the motor activity of DNA2 to displace RPA prior to resection. Using ensemble and single molecule biochemistry, we show that phosphorylated CtIP dramatically stimulates the ATP hydrolysis driven motor activity of DNA2. This activation in turn strongly promotes the degradation of RPA-coated ssDNA by DNA2. The domains of CtIP required to stimulate DNA2 are separable from those that regulate the MRN complex. These results establish that CtIP couples both MRE11-dependent short and DNA2-dependent long-range resection, and show how the motor activity of DNA2 promotes resection. Our data explain the less severe resection defects of MRE11 nuclease-deficient cells compared to those lacking CtIP.

**Highlights:**

- Phosphorylated CtIP stimulates the motor activity of DNA2
- The activated DNA2 translocase facilitates degradation of RPA-coated ssDNA
- CtIP promotes both MRN and DNA2 nucleases coupling short and long-range resection
- The CtIP domains required to promote DNA2 and MRN are distinct and fully separable

## Introduction

DNA double-strand breaks (DSBs) in eukaryotes are repaired by end-joining or homology-directed pathways (Ranjha et al., 2018). End-joining, including both non-homologous and microhomology-mediated sub-pathways require only minimal processing of the DNA ends and may involve limited microhomology at the break sites to facilitate ligation. Homologous recombination instead utilizes genetic information stored in an intact DNA copy, usually the sister chromatid in vegetative cells, to serve as a template for mostly accurate repair. There are several distinct sub-pathways of homology-directed repair, including single-strand annealing that leads to large deletions, synthesis-dependent strand annealing, break induced replication and canonical DNA double-strand break repair (Ranjha et al., 2018). While these pathways differ in mechanisms and resulting genetic outcomes, their common denominator is the first step, termed DNA end resection. All recombination pathways are initiated by nucleolytic degradation of the 5’-terminated DNA strand at DSB sites, leading to extended stretches of 3’-terminated ssDNA, which are required for the downstream steps in all the respective pathways (Bonetti et al., 2018; Cejka, 2015). Extensive DNA end resection commits the DSB repair to recombination, as resected ends are no longer ligatable by end-joining pathways. Cyclin-dependent kinases (CDK) regulate DNA end resection to be allowed only in the S and G2 phases of the cell cycle, when sister chromatids are available (Aylon et al., 2004; Cannavo et al., 2018; Chen et al., 2011; Huertas et al., 2008; Huertas and Jackson, 2009; Ira et al., 2004).

Intensive research established that DNA end resection generally consists of two subsequent steps. In human cells, resection is initiated by the MRE11-RAD50-NBS1 (MRN) complex, which functions in conjunction with CtIP (Sartori et al., 2007; Shibata et al., 2014). CtIP activates the MRE11 endonuclease within the MRN complex (Anand et al., 2016; Cannavo and Cejka, 2014). The activated MRN complex then preferentially cleaves the 5’-terminated DNA strand past protein blocks. Therefore, this step is particularly important to process DNA ends with non-canonical structures, including bound proteins like Ku and topoisomerases, as well as secondary DNA structures (Hoa et al., 2016; Liao et al., 2016; Reginato et al., 2018; Wang et al., 2017; Wilkinson et al., 2019). CtIP is phosphorylated by multiple kinases including CDK, ATM and ATR. In particular, the T847 CDK site of CtIP must be phosphorylated to promote MRN, which represents a mechanism that allows cell-cycle dependent control of DNA end resection (Anand et al., 2016; Huertas and Jackson, 2009; Liu et al., 2019; Orthwein et al., 2015; Polato et al., 2014).

Downstream of this initial short-range processing, either of two partially redundant nucleases catalyze long-range 5’-end resection that can extend up to several kilobases in length (Cannavo et al., 2013; Gravel et al., 2008; Liao et al., 2011; Mimitou and Symington, 2008; Nimonkar et al., 2011; Zhu et al., 2008). The Exonuclease 1 (EXO1) degrades 5’-terminated DNA strand within dsDNA. The nuclease-helicase DNA2 instead degrades ssDNA, and thus requires a helicase partner to unwind dsDNA, which can be either the Bloom (BLM) or Werner (WRN) RecQ family helicase (Pinto et al., 2016; Sturzenegger et al., 2014; Yan et al., 2011). Both BLM/WRN and DNA2 function in conjunction with the single-strand DNA binding protein RPA, which promotes DNA unwinding by BLM/WRN and directs DNA2 to degrade the 5’-terminated ssDNA strand (Cejka et al., 2010; Niu et al., 2010; Zhou et al., 2015). The structure of murine DNA2 revealed that the polypeptide possesses a central cavity through which ssDNA needs to thread before it can be cleaved (Zhou et al., 2015). As the size of the cavity only accommodates ssDNA, bound proteins, including RPA, need to be displaced prior to degradation (Balakrishnan et al., 2010; Zhou et al., 2015). The nuclease activity of DNA2 is essential for its function in resection and cell viability (Zhu et al., 2008). However, DNA2 also has a conserved helicase domain (Balakrishnan et al., 2010; Masuda-Sasa et al., 2006). Although the helicase-deficient DNA2 mutants are equally lethal, the function of this activity has remained enigmatic (Duxin et al., 2012). The helicase of DNA2 in resection cannot replace the helicase of BLM/WRN (Nimonkar et al., 2011). The unwinding capacity of wild type DNA2 is very weak, and processive DNA unwinding is only observed upon inactivation of the nuclease activity (Levikova et al., 2013; Masuda-Sasa et al., 2006; Pinto et al., 2016). The motor activity of DNA2 was thus proposed to promote resection not as a helicase to unwind dsDNA, but rather as a ssDNA translocase to act downstream of the RecQ family helicase partner (Levikova et al., 2017; Miller et al., 2017). In this model, BLM and WRN function as the lead helicases that unwind dsDNA. This provides long stretches of RPA coated 5’-terminated ssDNA to DNA2, which uses its motor activity to translocate along the ssDNA to displace RPA and feed the ssDNA strand to the nuclease domain (Levikova et al., 2017).

The phenotypes of human CtIP-depleted cells or corresponding *S. cerevisiae sae2*Δ mutants were reported to be more severe than those of MRE11/Mre11 nuclease point mutants, indicating that CtIP/Sae2 have additional functions that go beyond stimulating the MRE11 nuclease. To this point, both Sae2 and CtIP were described to possess an intrinsic nuclease activity with functions in DNA end resection and beyond, which could explain the phenotypic differences (Lengsfeld et al., 2007; Makharashvili et al., 2014; Wang et al., 2014). However, the Sae2/CtIP nuclease function remains controversial (Andres and Williams, 2017; Cannavo and Cejka, 2014; Wilkinson et al., 2019). Sae2/CtIP may also have a structural role in DNA break repair to bridge DNA (Andres et al., 2019; Wilkinson et al., 2019). In yeast, *sae2*Δ mutants were found to hyperactivate checkpoint signaling. Checkpoint-defective *sae2*Δ mutants exhibited identical DNA damage sensitivity and resection defects as *mre11* nuclease-dead mutants, showing that checkpoint hyperactivation accounted for the differential phenotypes of the respective mutants (Colombo et al., 2019; Gobbini et al., 2015; Yu et al., 2018). In higher eukaryotes, however, the situation is more complex. While some studies reported similar sensitivities of MRE11 nuclease-deficient (resulting from point mutations or small molecule inhibitors) and CtIP-deficient cells (Shibata et al., 2014), other reports point out more severe defects upon CtIP depletion. Specifically, in *Xenopus* egg extracts, complete inhibition of resection was observed upon depletion of EXO1 and CtIP, inferring that DNA2 requires CtIP to perform long-range resection (Peterson et al., 2013). Likewise, epistatic relationships between CtIP and DNA2 were found in chicken DT40 cells (Hoa et al., 2015b). Disruption of the MRE11 nuclease active site did not dramatically reduce resection, in contrast to CtIP-deficient cells, which exhibited a severe defect (Hoa et al., 2015a). Similarly, inactivation of the MRE11 nuclease was less severe than disruption of CtIP in human TK6 cells (Hoa et al., 2015a). Finally, single molecule analysis of DNA resection tracks showed that CtIP contributes to fast resection at long distances from the DNA end (Cruz-Garcia et al., 2014), which disagrees with the expected range of the MRN-dependent short-range resection. Together, these reports suggest that CtIP may promote long-range DNA end resection by an unknown mechanism. In accord, CtIP was shown to physically interact with and promote DNA unwinding by the BLM helicase, as well as to modestly stimulate DNA degradation by the DNA2 nuclease (Daley et al., 2017). Using ensemble and single molecule biochemistry, we show here that phosphorylated CtIP dramatically stimulates the motor activity of DNA2. This accelerates degradation of RPA-coated ssDNA by DNA2, showing the need for the motor activity of DNA2 to facilitate resection. Our results show that CtIP is thus a co-factor not only of MRE11, but also of DNA2, and demonstrate that the domains of CtIP required for the stimulation of MRN and DNA2 are physically separate. Our data support a model where CtIP first activates the MRE11 nuclease, and then helps couple short-range resection with the downstream long-range step by promoting DNA2. These results explain the dramatic DNA end resection defect observed in CtIP-depleted human cells.

## Results

### CtIP promotes DNA2-dependent long-range DNA end resection pathway

CtIP functions as a co-factor of the MRN endonuclease acting in the initial short-range DNA end resection pathway (Anand et al., 2016)(Fig. S1A). Multiple studies however indicated that deficiency of CtIP has a stronger impact on resection than mutations or inhibition of the MRE11 nuclease (Hoa et al., 2015a; Hoa et al., 2015b; Peterson et al., 2013) (see also Fig. S1B), suggesting that CtIP has additional functions in resection beyond promoting MRE11. To study the effect of CtIP on long-range DNA end resection pathways in a defined system, we expressed and purified phosphorylated human CtIP (pCtIP) in the presence of phosphatase inhibitors, DNA2, BLM, WRN and EXO1 (Fig. S1C–S1G) in *Sf*9 cells. Recombinant pCtIP did not promote DNA degradation by EXO1 (Fig. S1H), but stimulated DNA end resection by BLM-DNA2-RPA (Fig. S1I–S1K), as well as by WRN-DNA2-RPA (Fig. 1A, compare lanes 6 and 9). To understand the stimulation of resection by pCtIP in detail, we next investigated its effect on the activities of BLM/WRN and DNA2 individually. pCtIP stimulated DNA unwinding by BLM **~**2-fold (Fig. S1L), as well as to a similar extent the nuclease of DNA2 on oligonucleotide-based substrates (Fig. 1B–1C), as noted previously (Daley et al., 2017). In contrast, pCtIP did not significantly promote DNA unwinding by WRN (Fig. S1M). The strongest stimulatory effect by pCtIP was observed when we assayed DNA unwinding by nuclease-dead DNA2 D277A (Fig. 1D–1E). With pCtIP, 0.25 nM DNA2 D277A unwound more than 50% of the Y-structured DNA substrate, which was more than that unwound by 30 nM DNA2 D277A without pCtIP, corresponding to **>**10-fold stimulation (Fig. 1D–1E). Recombinant pCtIP thus predominantly promotes the motor activity of DNA2.

**Figure 1.**
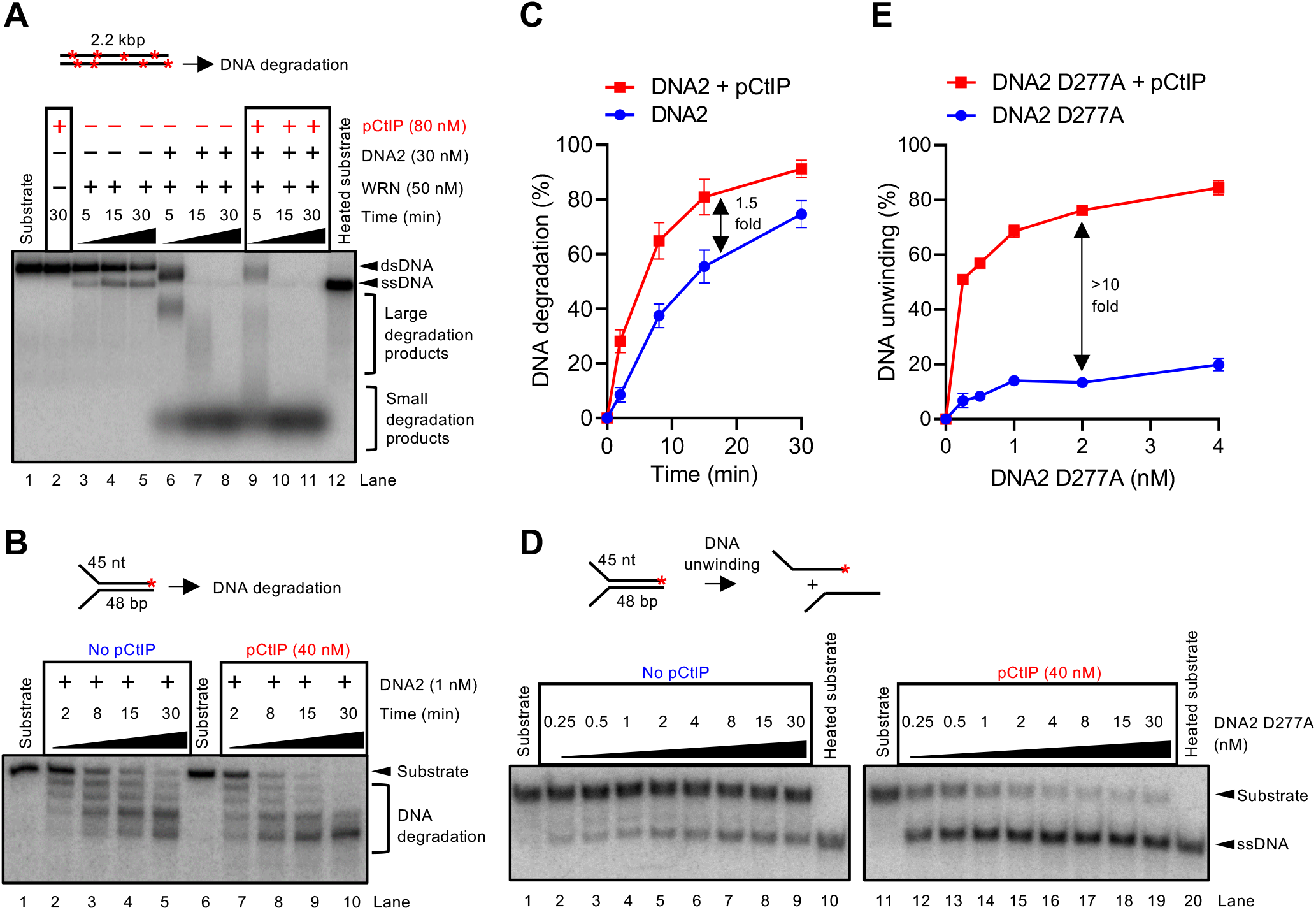
pCtIP promotes DNA2-dependent long-range DNA end resection pathway. A. DNA end resection by WRN, DNA2 and human RPA (176 nM) using 2.2 kbp-long randomly labelled dsDNA substrate in the presence or absence of phosphorylated CtIP (pCtIP). The reaction buffer contained 50 mM NaCl. Reaction products were separated by 1% agarose gel electrophoresis. Panel shows a representative experiment. Red asterisks indicate random labelling. B. Representative 15% denaturing polyacrylamide gel showing the degradation kinetics of a Y-structured (45nt/48 bp) DNA by DNA2 with or without pCtIP, in the presence of human RPA (15 nM) and 100 mM NaCl. Red asterisk indicates the position of the labelling. C. Quantitation of overall substrate utilization from experiments such as shown in panel B. N=3; error bars, SEM. D. A representative experiment showing DNA unwinding by nuclease-deficient DNA2 D277A with or without pCtIP on oligonucleotide-based Y-structured (45 nt/48 bp) DNA. Reactions were supplemented with human RPA (7.5 nM) and 50 mM NaCl and analysed on 10% native acrylamide gel electrophoresis. Red asterisk indicates the position of the labelling. E. Quantitation of overall substrate unwinding from experiments such as shown in panel D. N=3; error bars, SEM.

### CtIP dramatically promotes long-range ssDNA degradation by DNA2

To define the function of pCtIP in regulating DNA2 in a simple system, we next used an assay that monitors 5’→3’ degradation of 3’ end-labeled fragments of ssDNA by wild type DNA2. The use of ssDNA bypasses the requirement for BLM or WRN helicase. Without pCtIP, the degradation of ssDNA by DNA2 was slow (Fig 2A, lanes 3-7), and was strikingly **~**10-fold stimulated when pCtIP was included in the reactions (Fig 2A, lanes 9-13, Fig. 2B). While pCtIP appears to accelerate ssDNA degradation by DNA2, it did not change the 5’→3’ polarity of DNA degradation in the presence of RPA (Fig. S2A–S2B). Likewise, pCtIP did not allow DNA2 to cleave ssDNA endonucleolytically when DNA ends were blocked (Fig. S2A–S2B). Low nanomolar concentrations of both pCtIP and DNA2 were required for maximal DNA degradation under our conditions (Fig. 2C–2F). We note that 1 nM DNA2 in the presence of pCtIP was more efficient in DNA degradation than 20 nM DNA2 without pCtIP (compare Fig 2E, lane 7 with Fig. 2A, lane 7), highlighting the dramatic stimulation of the ssDNA degradative capacity of DNA2 by pCtIP.

**Figure 2.**
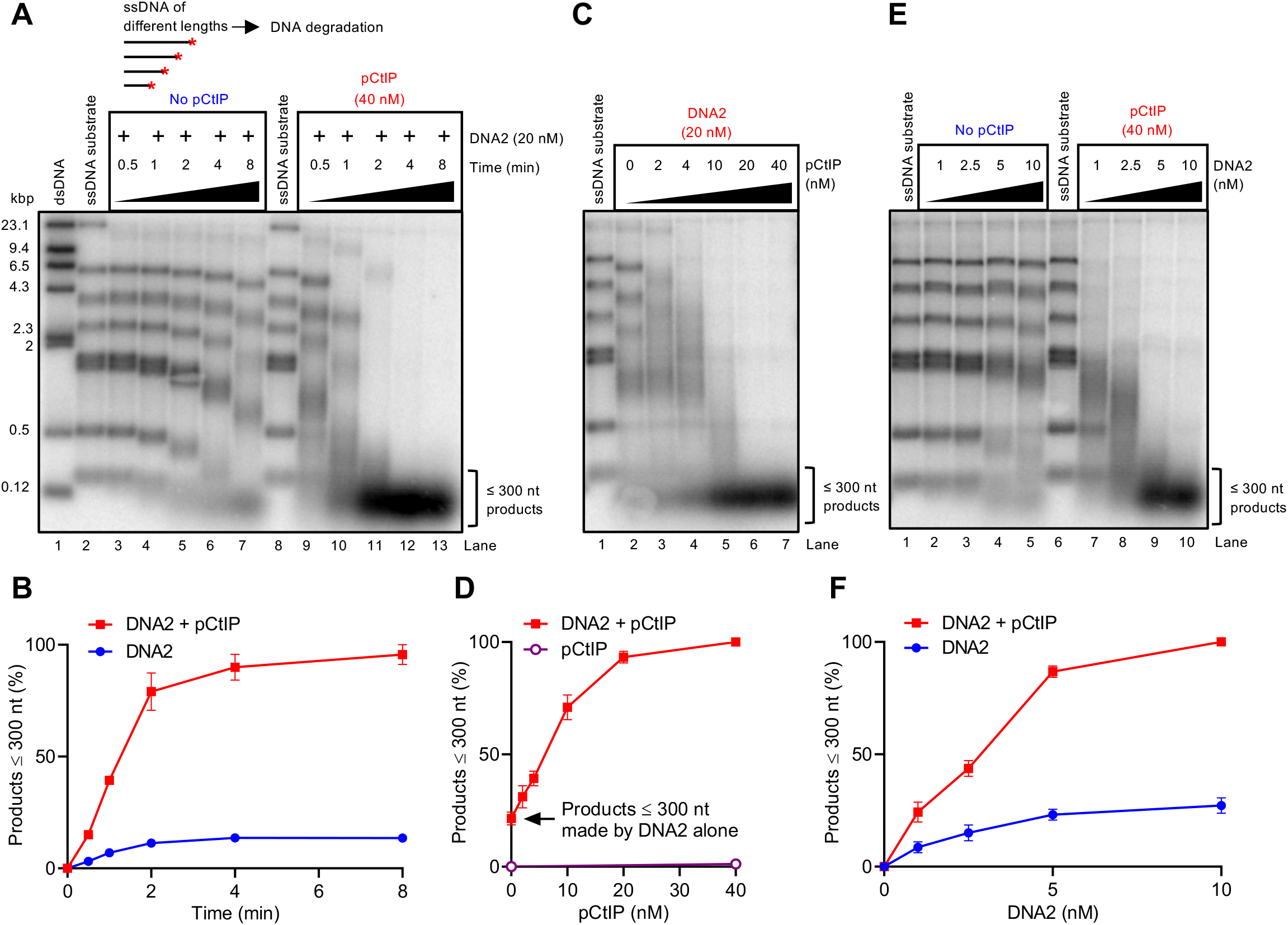
pCtIP dramatically promotes degradation of long stretches of ssDNA by DNA2. A. Representative 1% agarose gel showing degradation kinetics of 3’ ^32^P-labeled ssDNA fragments (derived from λDNA) of various lengths by DNA2 without or with pCtIP in the presence of 864 nM human RPA. The sizes of the corresponding dsDNA fragments are indicated on the left. Red asterisk indicates the position of the labelling. B. Quantitation of products smaller than **~**300 nt from experiments such as shown in panel A. N=3; error bars, SEM. C. Representative experiment as in A with various concentration of pCtIP incubated for 8 min. D. Quantitation of data such as shown in panel C; N=3; error bars, SEM. The degradation activity of pCtIP alone is the same as in Fig. 3A (lane 7). E. Experiment such as in A with various concentration of DNA2 incubated for 8 min in the presence or absence of pCtIP. F. Quantitation of data such as shown in panel E; N=3; error bars, SEM.

**Figure 3.**
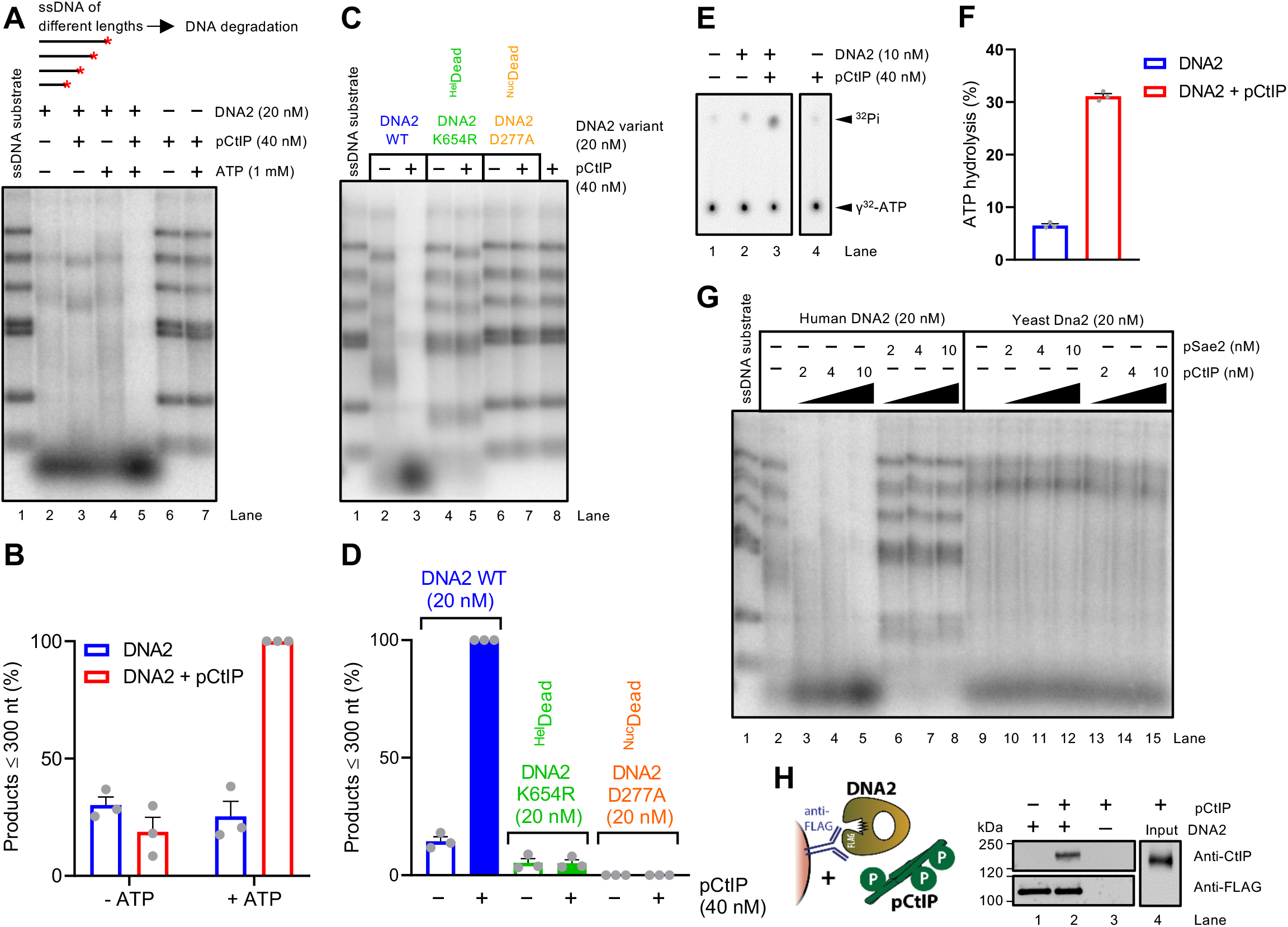
The motor activity of DNA2 mediates the accelerated ssDNA degradation with pCtIP. A. Degradation of ssDNA fragments (derived from λDNA) of various lengths by DNA2 in the presence or absence of pCtIP without or with ATP, as indicated. All reactions contained human RPA (864 nM). The experiment was incubated at 37°C for 8 min. ATP is required for the stimulatory effect of pCtIP on DNA2. Red asterisk indicates the position of the labelling. B. Quantitation of data such as shown in panel A. N=3; error bars, SEM. C. Degradation of ssDNA fragments by wild type, helicase-deficient K654R or nuclease-deficient D277A DNA2 variants without or with pCtIP. All reactions contained human RPA (864 nM). The experiment was incubated at 37°C for 8 min. D. Quantitation of small degradation products from experiment such as shown in panel C. N=3; error bars, SEM. E. ATP hydrolysis by DNA2 alone (10 nM) or with pCtIP (40 nM). Reactions contained 10.3 kbp-long substrate denatured at 95°C for 5 min, 395.5 nM human RPA and no added salt. F. Quantitation of ATP hydrolysis from experiments such as shown in panel E. N=3; error bars, SEM. G. Degradation of ssDNA fragments by human or yeast DNA2/Dna2 without or with human pCtIP or yeast pSae2. Reactions with human DNA2 were carried out at 37°C for 8 min with human RPA (864 nM). Reactions with yeast Dna2 were performed at 25°C for 1 min with yeast RPA (1.09 µM). H. Analysis of DNA2 interaction with pCtIP. DNA2-FLAG was immobilized on M2 anti-FLAG affinity resin and incubated with purified recombinant pCtIP. The Western blot was performed with anti-FLAG and anti-CtIP antibodies.

### The motor activity of DNA2 mediates the accelerated ssDNA degradation with pCtIP

To define the mechanism of the stimulatory effect of pCtIP on the DNA2 nuclease, we set out to investigate the role of the motor activity of DNA2 in this process. Strikingly, pCtIP strongly stimulated ssDNA degradation by DNA2 only in the presence of ATP (Fig. 3A–3B). Importantly, pCtIP had no activity on its own with or without ATP (Fig. 3A, lanes 6 and 7). Furthermore, pCtIP did not notably promote ssDNA degradation by the helicase-deficient with dirsrupted ATPase site DNA2 K654R variant (Fig. 3C–3D). We also note that pCtIP strongly promoted the ATPase activity of DNA2 (Fig. 3E–3F), while no ATP hydrolysis was observed when using the DNA2 K654R variant, as expected (Fig. S3A). Assuming our DNA2 preparation is 100% active, the experiments indicate an apparent *k*_*cat*_ for DNA2 without pCtIP of **~**10 sec^−1^, which was **~**5-fold stimulated when pCtIP was included in the reaction. These experiments together demonstrate the importance of ATPase-driven translocase activity of DNA2 in extended ssDNA degradation.

No DNA degradation was observed when using the nuclease-deficient DNA2 D277A (Fig. 3C–3D) even in the presence of pCtIP, further demonstrating that the observed nuclease activity was intrinsic to DNA2 and not a result of non-specific contamination in our assays. Human pCtIP did not promote ssDNA degradation by yeast Dna2 and *vice versa*, yeast pSae2 did not promote the nuclease of human DNA2 (Fig. 3G, Fig. S3B–S3C), implicating direct species-specific interactions between the human polypeptides. Furthermore, pSae2 did not notably stimulate yeast Dna2, showing that the interplay described here is specific to higher eukaryotes. The species-specific interplay of cognate human pCtIP and DNA2 suggested the proteins might directly physically interact. Indeed, FLAG-tagged DNA2 could readily pull down pCtIP, demonstrating a direct interaction (Fig. 3H). Together, these results establish that by forming a complex, pCtIP stimulates the motor activity of DNA2, which in turn accelerates its ssDNA degradative capacity.

### Single-molecule experiments reveal the acceleration of DNA unwinding rate of DNA2 by pCtIP

To further mechanistically define the stimulation of DNA2 motor by pCtIP, we employed magnetic tweezers, which monitor the position of a magnetic bead linked by a single DNA molecule to a fixed surface. As ssDNA is extended compared to dsDNA, DNA unwinding can be inferred from the relative position of the bead and DNA extension curve. We could not directly monitor the DNA2 nuclease in our setup, because DNA must be attached on both ends. Instead, we analyzed the effect of pCtIP on DNA unwinding by the nuclease-dead DNA2 D277A (Fig. 4A). The mean velocity of DNA2 D277A in the absence of pCtIP was 13 ± 1 bp/s (SEM), which was increased **~**7-fold in the presence of pCtIP to 86 ± 4 bp/s (SEM)(Fig. 4B–C and Fig. S4A). In contrast, pCtIP had no effect on the processivity of DNA2 D277A (Fig. S4B). While the motor activity of human DNA2 alone is slower than that of the yeast orthologue (Levikova et al., 2013; Pinto et al., 2016), we note that in the presence of pCtIP, the unwinding activity of human DNA2 is faster than that by yeast Dna2, making human DNA2 one of the most efficient known motor proteins in eukaryotes.

**Figure 4.**
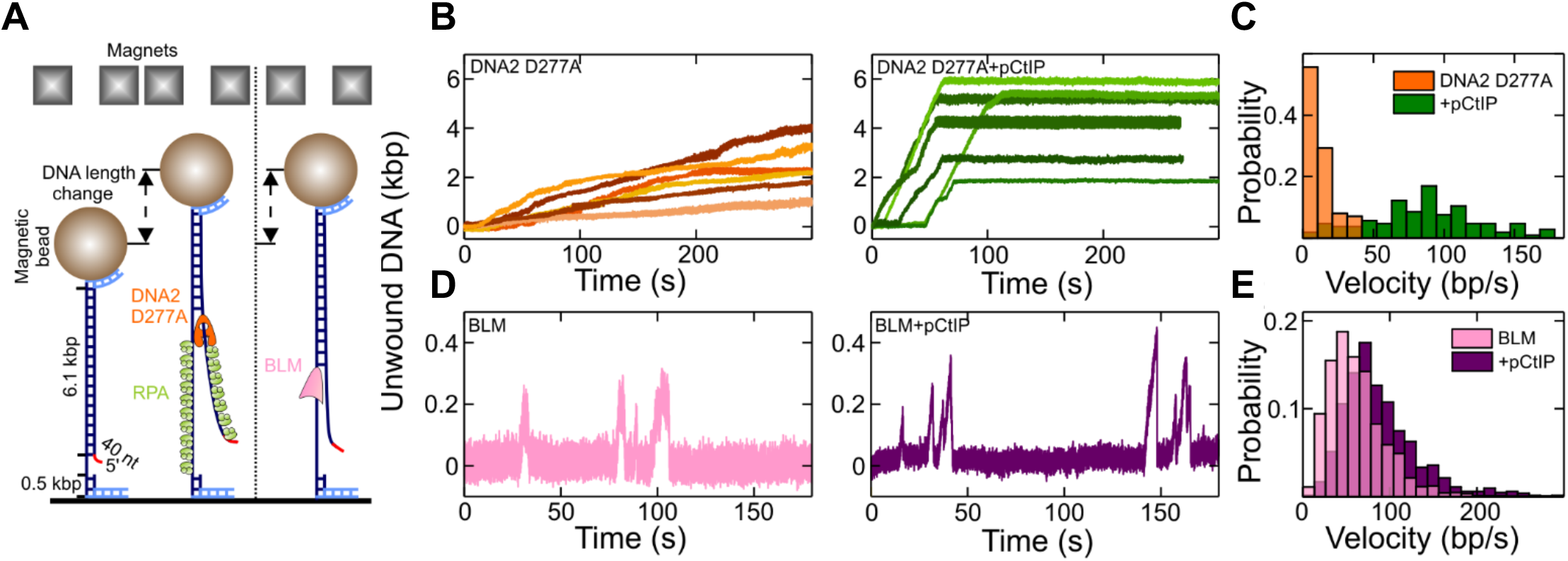
Single-molecule experiments demonstrate accelerated DNA2 motor activity in the presence of pCtIP. A. Sketch of the employed magnetic tweezers assay and the DNA construct carrying a 40 nt 5’ flap to allow loading of either helicase. B. Representative DNA unwinding events of the DNA2 nuclease-dead mutant (D277A, 25 nM) in the absence (orange) and presence (green) of pCtIP (25 nM). Both reactions were also supplemented with 25 nM human RPA. C. Histograms of the observed unwinding velocities for nuclease-dead DNA2 (N = 40 traces for each case). D. Representative DNA unwinding events of BLM (25 nM) in the absence (magenta) and presence (purple) of pCtIP (25 nM). E. Histogram of the observed unwinding velocities for BLM. (N = 852 events). The force in all experiments was F = 19 ± 3 pN (SD).

As pCtIP also modestly promotes DNA unwinding by BLM (Fig. S1L)(Daley et al., 2017), we also assayed this stimulatory effect by magnetic tweezers. The DNA unwinding by BLM, as by other helicases from the RecQ family such as Sgs1, is characterized by bursts of DNA unwinding followed by rewinding (Fig. 4D)(Kasaciunaite et al., 2019). We observed **~**1.5-fold stimulation of DNA unwinding by BLM when pCtIP was included in the reaction (Fig. 4D–E). The mean BLM velocity in the absence of pCtIP was 61 ± 4 bp/s (SEM), and 91 ± 4 bp/s (SEM) in the presence of pCtIP. As with DNA2 D277A, pCtIP did not affect the processivity of BLM (Fig. S4C). Additionally, pCtIP did not stimulate DNA unwinding by WRN (Fig. S4D–F), in agreement with the biochemical data (Fig. S1M). No DNA unwinding was observed without ATP, or by CtIP alone (Fig.S4G–K). In summary, pCtIP promotes the velocity of DNA unwinding by both BLM and DNA2 without affecting their processivity. The effect of pCtIP on DNA2 motor is much greater than that on BLM (Compare Fig. 4C and Fig. 4E).

### Domains of pCtIP that promote MRN and DNA2 are fully separable

The pCtIP protein is composed of an N-terminal tetramerization domain, an unstructured central region - largely lacking in *S. cerevisiae* -that binds DNA and contains a number of phosphorylation sites, and finally a conserved C-terminal domain that is required to activate MRN, has a secondary DNA binding site and is likewise subject to phosphorylation (Fig. 5A)(Anand et al., 2016; Davies et al., 2015; Sartori et al., 2007; Wilkinson et al., 2019). A key cyclin-dependent kinase site, T847, which enables cell-cycle regulated control of DNA end resection by licensing stimulation of MRN, is located within the C-terminal domain of pCtIP (Fig. 5A) (Huertas and Jackson, 2009). Only the N-terminal tetramerization and the C-terminal domains bear limited similarity to yeast Sae2. To identify pCtIP regions that are required for the stimulation of DNA2, we designed pCtIP variants lacking stretches of amino acids from the internal region such as pCtIP Δ1 (lacking residues 350-600 including the DNA-binding region) and pCtIP Δ2 (lacking residues 165-790 comprising the entire region between the tetramerization domain and the Sae2-like C-terminal domain). As wild type full-length pCtIP, these internally truncated variants were expressed in *Sf*9 cells and purified in the presence of phosphatase inhibitors to preserve phosphorylation. We observed that pCtIP Δ1, but not pCtIP Δ2, was fully proficient in stimulating DNA2 (Fig. 5B–5C, Fig. S5A). This indicated that the region in pCtIP between residues 350 and 600 containing the DNA binding domain is dispensable for the stimulation of DNA2, and the stimulatory activity is dependent on a region upstream or downstream of these residues. We also prepared the C-terminal domain of pCtIP alone (pCtIP Δ3, containing residues 790-897), and a mutant lacking this C-terminal domain (pCtIP Δ4) (Fig. 5A). Remarkably, the pCtIP Δ4 variant, lacking the C-terminal Sae2-like region, while incapable to stimulate MRN (Fig. S5B), as expected (Anand et al., 2016), was fully proficient in stimulating DNA2 (Fig. 5B–5C, Fig. S5A). In accord with these data, pCtIP Δ4, but not the pCtIP Δ2 variant, was capable to promote the ATPase of DNA2 (Fig. S5C–F). Therefore, the regions of pCtIP that are required to promote MRN and DNA2 are distinct and physically separate.

**Figure 5.**
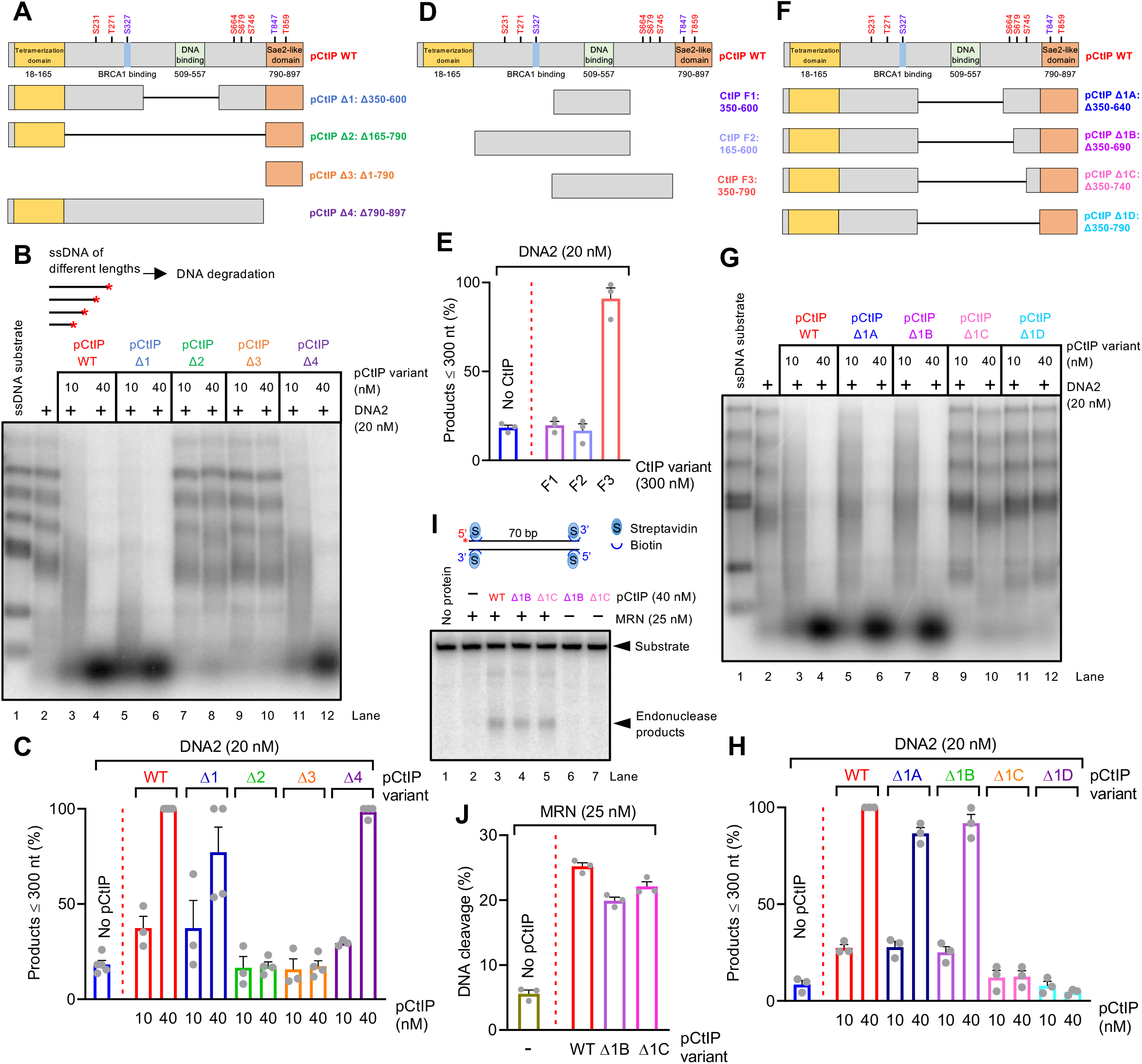
Separate domains of pCtIP promote the MRN and DNA2 nucleases. A. A schematic representation of the primary structure of wild type pCtIP and internal deletion variants (pCtIP Δ1-4) purified from *Sf*9 cells in the presence of phosphatase inhibitors. Main ATM phosphorylation sites are indicated in red, CDK phosphorylation sites are indicated in blue. B. Degradation ssDNA fragments of various lengths by DNA2 without or with pCtIP variants, as indicated. All reactions contained human RPA (864 nM). Red asterisk indicates the position of the labelling. C. Quantitation of data such as shown in panel B. N=3-4; error bars, SEM. D. A schematic representation of purified recombinant CtIP fragments (F1-F3) expressed in *E. coli*. pCtIP wild type purified from *Sf*9 cells is again shown as a reference. E. Quantitation of ssDNA fragment degradation into products smaller than **~**300 nt by DNA2 without or with F1-F3 CtIP fragments. N=3; error bars, SEM. F. A schematic representation of internal deletion variants (pCtIP Δ1A-Δ1D) purified from *Sf*9 cells in the presence of phosphatase inhibitors. pCtIP wild type is again shown as a reference. G. Degradation of ssDNA fragments of various lengths by DNA2 without or with various concentrations of wild type pCtIP or Δ1A-Δ1D mutants, in the presence of human RPA (864 nM). H. Quantitation of data such as shown in panel G. N=3; error bars, SEM. I. Endonuclease assay with MRN (25 nM) and wild type full-length or pCtIP Δ1B and Δ1C variants. Red asterisk indicates the position of the labelling. J. Quantitation of data such as shown in panel I. N=3; error bars, SEM.

To further map the CtIP region required to stimulate DNA2, we expressed various portions of the central CtIP domain in *E. coli* (Fig. 5D). We prepared the central region corresponding to residues 350-600, as well as fragments containing additionally the upstream and downstream regions from the wider central domain (Fig. 5D). We observed that CtIP F3 fragment, containing residues 350-790, was capable to promote DNA2 (Fig. 5E). This indicated that the CtIP region downstream of residue 600 was required for DNA2 stimulation. We note that this domain is lacking in yeast Sae2, and our result is thus in agreement with the observation that yeast Sae2 does not promote yeast or human DNA2 (Fig. 3G). We observed that a high concentration of the F3 fragment compared to pCtIP variants expressed in insect cells was needed to observe DNA2 stimulation (compare Fig. 5C and Fig. 5E), indicating a lower stimulatory capacity of the polypeptide expressed in *E. coli* compared to variants prepared in *Sf*9 cells.

To narrow down the stimulatory region in pCtIP even further, we expressed in *Sf*9 cells and purified an additional series of internal pCtIP truncations lacking also residues downstream of position 600 (Fig. 5F, Fig. S5G). We observed that pCtIP Δ1B fragment (lacking residues 350-690) was fully proficient in stimulating DNA2, but additional shortening of this mutant by 50 residues (pCtIP Δ1C, lacking residues 350-740) entirely eliminated this capacity (Fig. 5G–H, Fig. S5H). In contrast, the pCtIP Δ1C construct was fully proficient in stimulating the endonuclease of MRN (Fig. 5I-J). These experiments established that the pCtIP region immediately downstream of residue 690 is required for the stimulation of the DNA2 motor activity, while it is dispensable for the stimulation of MRN.

We also note that pCtIP NA/HA, which was described to lack intrinsic nuclease activity (Makharashvili et al., 2014), and was recently found deficient in its interplay with DNA2 in response to stalled replication forks (Przetocka et al., 2018), was partially impaired in both stimulating the MRN complex (Fig. S5I–S5L) as well as DNA2 (Fig. S5M–S5N). Although the NA/HA mutations lie outside of the region directly involved in the stimulation of DNA2 (Fig. S5J), they may affect the structure of the CtIP central domain and thus broadly impair CtIP functions. The NA/HA mutations may thus not strictly reflect an intrinsic nuclease defect. Finally, DNA2 nuclease and helicase-dead (DNA2 D277A/K654R) had no effect on the endonuclease activity of the MRN-pCtIP complex (Fig. S5O). Together, these results establish that pCtIP promotes MRN and DNA2 via distinct domains that are fully separable, and map the minimal region required for the stimulation of DNA2 between residues 690 and 790 of pCtIP.

### pCtIP phosphorylation facilitates its capacity to promote DNA2

Due to the established requirement for pCtIP phosphorylation in the stimulation of MRN *in vitro* (Anand et al., 2019; Anand et al., 2016), we set out to define the function of pCtIP phosphorylation in the regulation of DNA2. Dephosphorylation of full-length, Δ1 and Δ4 pCtIP variants expressed in *Sf*9 cells dramatically reduced their capacity to stimulate DNA2 (Fig. 6A–6B, Fig. S6A–S6C). Likewise, lambda phosphatase treatment of pCtIP reduced its capacity to promote DNA unwinding by DNA2 D277A (Fig. S6D–S6E). This is in agreement with data from Fig. 5E, where we observed limited activity of the CtIP fragment expressed in *E. coli*, which was expected not to be phosphorylated.

**Figure 6.**
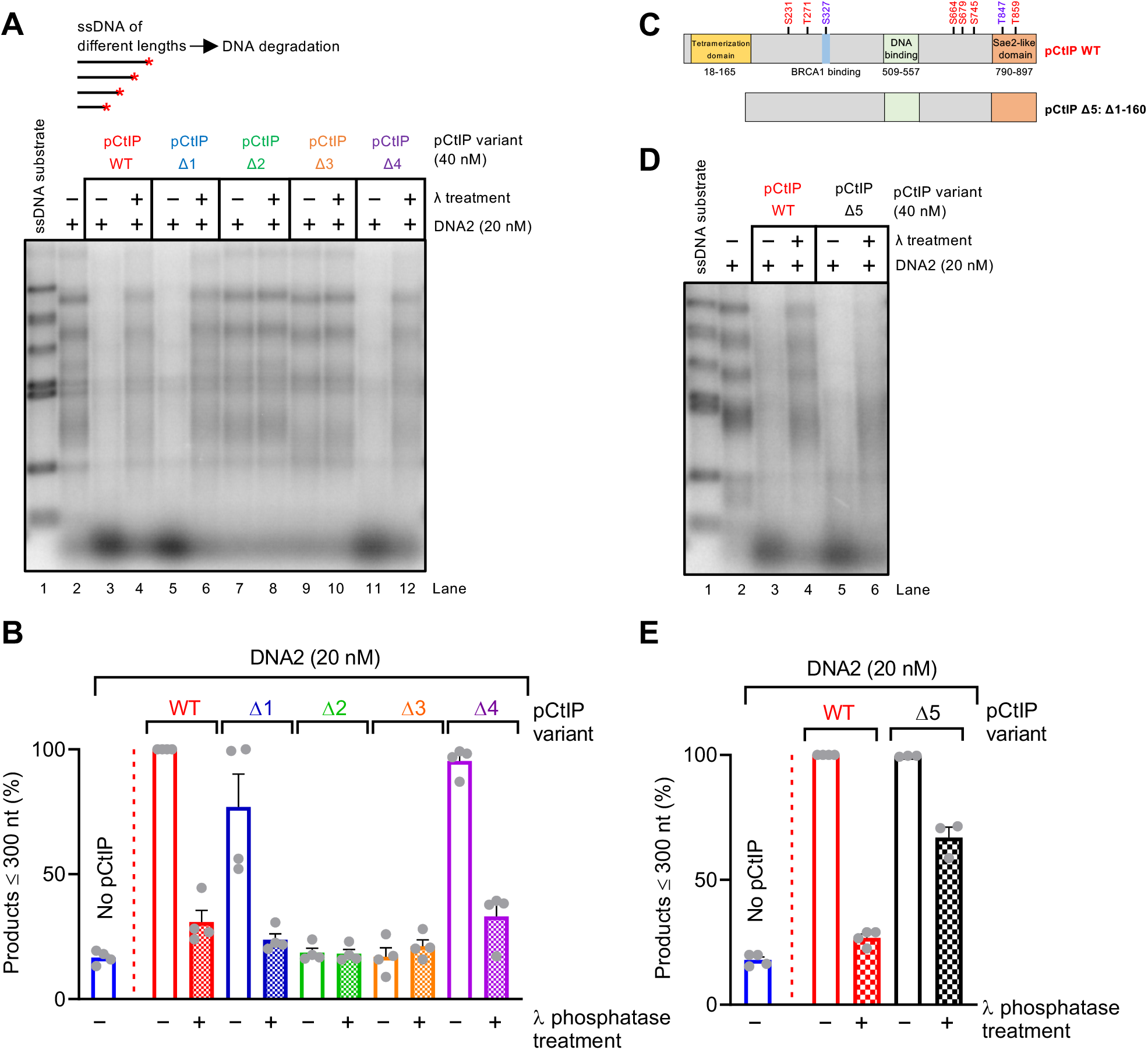
pCtIP phosphorylation facilitates its capacity to promote DNA2. A. Degradation of ssDNA fragments of various length with DNA2 alone (20 nM) or with pCtIP wild type, Δ1, Δ2, Δ3, Δ4, mock-treated or λ-treated, in the presence of human RPA (864 nM). Red asterisk indicates the position of the labelling. B. Quantitation of small degradation products from experiments such as shown in panel A. N=4; error bars, SEM. C. A schematic representation of purified recombinant pCtIP Δ5 lacking the first 160 amino acids. pCtIP wild type is again shown as a reference. D. Degradation of ssDNA fragments of various length with DNA2 alone (20 nM) or with pCtIP wild type or Δ5, mock-treated or λ-treated, in the presence of human RPA (864 nM). E. Quantitation of data such as shown in panel D. N=3-4; error bars, SEM.

Dephosphorylated CtIP and their homologs are thought to aggregate, so the apparent stimulatory function of phosphorylation may stem from abrogated CtIP aggregation (Cannavo et al., 2018; Fu et al., 2014). To address whether phosphorylation enables CtIP to promote DNA2 beyond preventing aggregation, we employed pCtIP Δ5, which lacks the N-terminal tetramerization domain (residues 1-160)(Andres et al., 2015; Davies et al., 2015)(Fig. 6C). Lack of the tetramerization domain is expected to prevent aggregation, even in the dephosphorylated state (Cannavo et al., 2018). This pCtIP variant was proficient in stimulating DNA2, and was partially resistant to lambda phosphatase treatment (Fig. 6D–6E, Fig. S6F–S6G). We then analyzed a number of point mutants non-phosphorylatable on some of the previously-established phosphorylation sites in pCtIP, including S664A, S679A, S754A and T847A (Fig. S6H). None of these mutations eliminated the capacity of pCtIP to promote DNA2 (Fig. S6I), indicating that these phosphorylation sites are either functionally redundant, or that phosphorylation on residues other than those analyzed is important for the observed activity.

We conclude that pCtIP phosphorylation clearly facilitates the stimulation of DNA2. This is likely caused, at least in part, indirectly by preventing pCtIP aggregation. Furthermore, while CtIP tetramerization is essential to stimulate MRN (Anand et al., 2016; Davies et al., 2015), it may be dispensable to stimulate DNA2.

## Discussion

CtIP has a well-established role as an activator of the MRE11 nuclease activity within the MRN complex, which initiates DNA end resection. This function is conserved in evolution from yeast to human cells (Anand et al., 2016; Cannavo and Cejka, 2014; Deshpande et al., 2016; Wang et al., 2017). The results presented here demonstrate that CtIP also controls DNA2, a long-range DNA end resection nuclease that functions downstream of MRE11. Therefore, similarly to SLX4 that controls a number of structure specific nucleases acting late in the recombination pathway (Fekairi et al., 2009; Wyatt et al., 2017), CtIP controls both short-range and long-range resection nucleases (Fig. 7A). We propose that coupling of both resection steps by CtIP allows to better coordinate and regulate DNA end resection. The CtIP domains required to promote MRE11 and DNA2 are at least partially separate. We found that the very N-terminal and C-terminal domains of CtIP, while being required to promote MRE11, are largely dispensable to regulate DNA2 (Fig. 7B). Instead, we identified a region in the central domain of pCtIP between residues 690 and 790 that is essential to stimulate DNA2, but dispensable for stimulating MRN (Fig. 7B). The central domain of CtIP is lacking in homologues of low eukaryotes such as in *S. cerevisiae* Sae2. In accord, we failed to observe stimulation of yeast Dna2 by yeast Sae2. Therefore, while the regulation of MRE11 by CtIP is conserved in evolution, the regulation of DNA2 appears to be restricted to higher eukaryotes.

**Figure 7.**
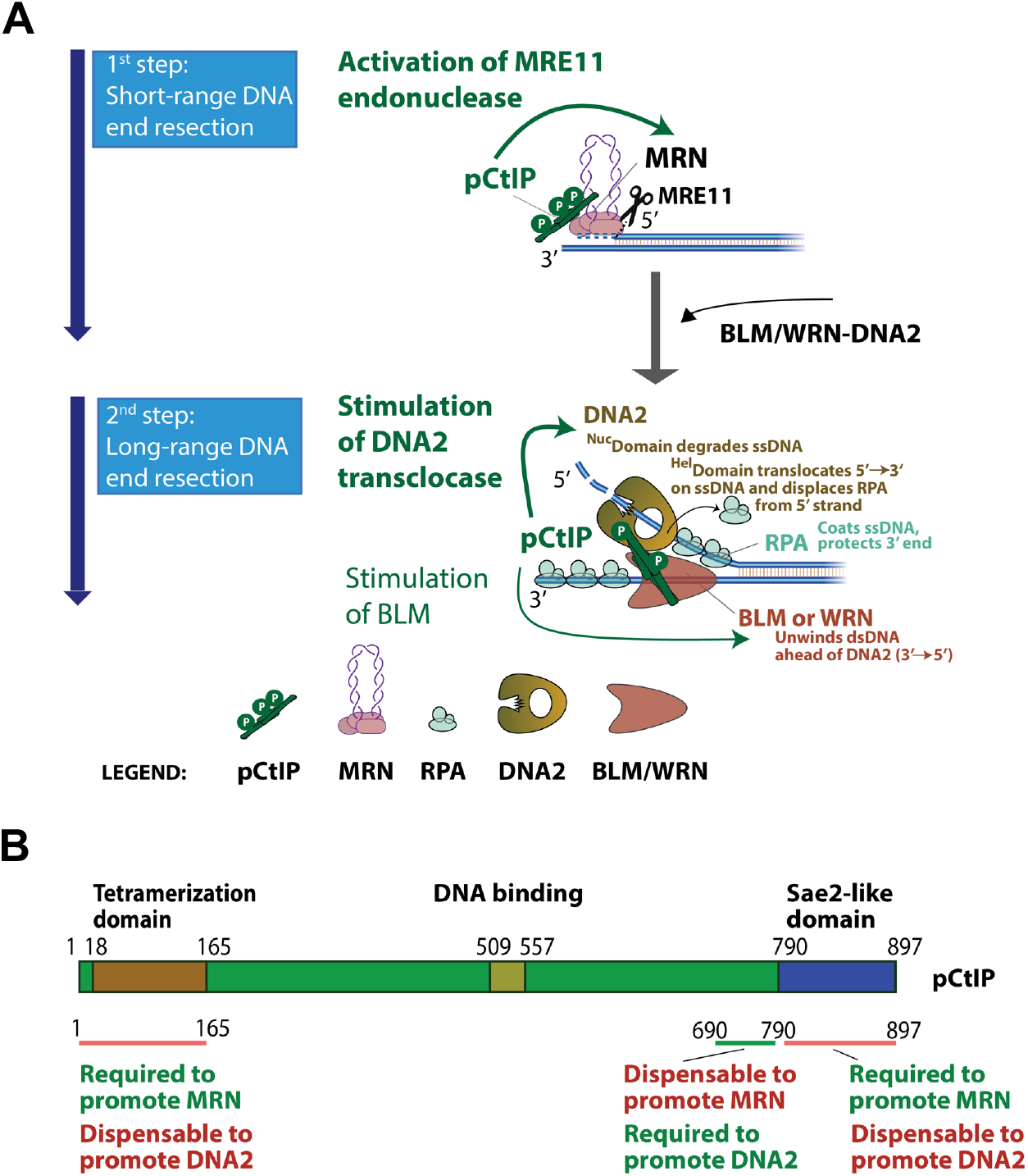
Model for pCtIP functions in DNA end resection. A. A cartoon depicting the role of pCtIP in short-range resection by MRE11 within the MRN complex and in long-range resection by DNA2-BLM. In long-range resection, pCtIP stimulates both the translocase activity of DNA2 to facilitate degradation of RPA-coated ssDNA, and the helicase activity of BLM to unwind dsDNA. B. A schematic representation of the primary structure of pCtIP and the domains required for the stimulation of MRN or DNA2 nucleases.

Deletion of *SAE2* results in only limited resection defects in yeast cells. While Sae2 and the Mre11 nuclease are critical to remove Spo11, stalled topoisomerases or secondary DNA structures, the MRX-Sae2 pathway is partially dispensable for the processing of clean DSBs in yeast (Llorente and Symington, 2004). In contrast, defects in CtIP dramatically inhibit DNA end resection in human cells, including that of nuclease-generated clean breaks (Sartori et al., 2007). Accordingly, depletion of CtIP is used to abrogate resection in many experimental systems in human cells. The regulation of DNA2 by CtIP presented here thus could help explaining the differential requirement for Sae2 and CtIP in resection in high and low eukaryotes. At the same time, it is possible that the short-range resection may be generally more important in human cells. The critical function of CtIP could thus also result from an upstream placement of the MRN-CtIP-dependent DNA clipping. Abrogation of a critical upstream step would then mask an involvement of CtIP downstream of MRE11, explaining similar phenotypes resulting from downregulation of CtIP and chemical inhibition of the MRE11 nuclease observed in some studies (Shibata et al., 2014). Other studies, including ours (Fig. S1B)(Hoa et al., 2015a; Hoa et al., 2015b), in contrast noted a more severe resection defects associated with CtIP depletion compared to inhibition or mutagenesis of the MRE11 nuclease active site, implicating other functions for CtIP beyond the regulation of MRE11. The differences may be due to the experimental systems used, which may have different requirements for the initial short-range resection by MRN-CtIP. Nevertheless, studies in human and murine cells, as well as *Xenopus* egg extracts, noted that CtIP may promote the long-range dependent pathway by DNA2 (Hoa et al., 2015a; Hoa et al., 2015b; Peterson et al., 2013). The results presented here provide a mechanistic explanation for the CtIP and DNA2 function in this process.

Our data indicate that CtIP primarily functions to stimulate the ATPase-dependent ssDNA translocase activity of DNA2. While DNA2 is primarily active as a nuclease, the function of its helicase domain remained somewhat elusive. We previously reported that the helicase activity of DNA2 becomes only apparent after inactivation of the nuclease (Levikova et al., 2013; Pinto et al., 2016). Therefore, DNA2 motor does not likely function as a helicase to unwind dsDNA, but rather as a ssDNA translocase, which is readily observed with the wild type protein (Levikova et al., 2017; Miller et al., 2017). According to this model, the ssDNA translocase activity helps feed ssDNA to the nuclease domain of DNA2 and facilitates thus its degradation. Moreover, DNA unwound by the cognate RecQ family helicase partner is coated by RPA, which directs the DNA2 nuclease to degrade the 5’-terminated ssDNA strand (Cejka et al., 2010; Niu et al., 2010). However, as only naked ssDNA can pass through the channel within the DNA2 structure to reach the nuclease domain (Zhou et al., 2015), RPA needs to be displaced, requiring an active mechanism. The motor activity of DNA2 plays an important function during this process, as it accelerates DNA degradation primarily when RPA is present (Levikova et al., 2017). The results presented here demonstrate that CtIP further accelerates the motor activity of DNA2, and thus indirectly stimulates ssDNA degradation by its nuclease domain (Fig. 7A). The stimulation of the DNA2 motor is not the only way how CtIP promotes the BLM-DNA2 long-range resection pathway. Sung and colleagues previously reported that CtIP also stimulates DNA unwinding by BLM (Daley et al., 2017), which functions upstream of DNA2 (Fig. 7A). We could confirm this stimulatory effect. In single molecule experiments, we observed that CtIP accelerated DNA unwinding by BLM by **~**1.5-fold, while it stimulated DNA2 by **~**10-fold under the same experimental conditions, demonstrating a comparatively larger effect of CtIP on DNA2 compared to BLM.

Finally, we demonstrate that phosphorylation of pCtIP facilitates its capacity to promote DNA2. Therefore, similarly to the interplay with MRE11, phosphorylation represents a key regulatory mechanism that keeps the DNA2 activity in check. Together, this allows resection only in the S-G2 phases of the cell cycle and upon DNA damage, which limits illegitimate recombination.

## Experimental procedures

### Cloning, expression and purification of recombinant proteins

Wild-type DNA2, nuclease-dead DNA2 D277A, helicase-dead DNA2 K654R and double-dead DNA2 D277A K654R (DNA2 DA/KR) were expressed in *Spodoptera frugiperda* 9 (*Sf*9) insect cells and purified by affinity chromatography exploiting the N-terminal 6X his-tag and the C-terminal FLAG-tag (Anand et al., 2018; Pinto et al., 2016). Yeast Dna2 was expressed in the *S. cerevisiae* strain WDH668 and purified using the N-terminal FLAG tag and the C-terminal 6X his-tag (Levikova et al., 2013). MRN was prepared using the 6X his and FLAG tags at the C-termini of MRE11 and RAD50, respectively (Anand et al., 2018; Anand et al., 2016). BLM, WRN, Sae2 and wild-type CtIP, CtIP Δ5 or CtIP T847A were obtained taking advantage of the maltose-binding protein (MBP) tag at the N-terminus and 10X his-tag at the C-terminus. The MBP tag was cleaved off during purification (Anand et al., 2016; Cannavo and Cejka, 2014; Cannavo et al., 2018; Pinto et al., 2016). The constructs for the expression of CtIP internal deletion mutants (Δ1 to Δ4 and Δ1A to Δ1D) were prepared by PCR to amplify and fused the corresponding CtIP sequences and cloned into NheI and XmaI sites of pFB-2XMBP-CtIP-10Xhis between the 2X MBP-tag and the 10X his-tag. CtIP N289A H290A (CtIP NA/HA) and CtIP non-phosphorylatable S664A, S679A and S745A variants were prepared by mutating the respective pFB-MBP-CtIP-his plasmid by QuickChange site-directed mutagenesis kit following manufacturer’s instructions (Agilent Technology). The truncated proteins and point mutants were expressed and purified using the same procedure as the full-length protein. For expression of phosphorylated CtIP (pCtIP) variants and phosphorylated Sae2, *Sf*9 cells were treated with 50 nM Okadaic acid (APExBIO) to preserve proteins in their phosphorylated state, and 1 µM camptothecin (Sigma) to further activate protein phosphorylation cascade (Anand et al., 2018; Anand et al., 2016). Where indicated, pCtIP was dephosphorylated with λ-phosphatase (New England Biolabs) according to the manufacturer’s instructions. For ‘‘mock’’ controls, λ-phosphatase was excluded from the reactions, and the sample was otherwise incubated in the same way as the λ-phosphatase-treated reaction. The constructs coding for the CtIP fragments F1 to F4 were fused with C-terminal 6X his-tag and cloned into BamHI and PstI sites in pMALT-P vector (Kowalczykowski laboratory) and expressed in *E. coli* BL21 DE3 pLysS cells. Purification was performed by affinity chromatography using amylose (New England Biolabs) and Ni-NTA (Qiagen) resins with the same buffers and procedure described for wild-type CtIP and its variants expressed in *Sf*9 cells (Anand et al., 2016). Human and yeast RPA were expressed in *E. coli* and purified using ÄKTA pure (GE Healthcare) with HiTrap Blue HP, HiTrap desalting and HiTrap Q chromatography columns (all GE Healthcare)(Anand et al., 2018). EXO1-FLAG was expressed in *Sf*9 cells and purified as described in detail in supplementary information. The sequence of all primers used for PCR and cloning in this study is listed in Supplementary Table 1.

### Preparation of DNA substrates

Oligonucleotide-based DNA substrates were ^32^P-labeled either at the 5ꞌ terminus with [γ-^32^P]ATP (Perkin Elmer) and T4 polynucleotide kinase (New England Biolabs), or at the 3 ꞌ terminus [α-^32^P]dCTP (Perkin Elmer) and terminal transferase (New England Biolabs) according to the manufacturer’s instructions (Pinto et al., 2018). Unincorporated nucleotides were removed using Micro Bio-Spin P-30 Tris chromatography columns (Biorad). To prepare the quadruple blocked 70 bp-long dsDNA substrate, the oligonucleotides PC210 and PC211 were used (Cannavo and Cejka, 2014). The sequence of all oligonucleotides used for DNA substrate preparation is listed in Supplementary Table 2. The Y-structured DNA substrate was prepared with the oligonucleotides X12-3HJ3 and X12-3TOPL (Pinto et al., 2016). The randomly labeled 2.2-kbp-long substrate was prepared amplifying the human NBS1 gene by PCR from pFB-MBP-NBS1-his plasmid (Anand et al., 2018) using Phusion high-fidelity DNA polymerase (New England Biolabs) and the NBS1_F and NBS1_R primers. 66 nM [α-^32^P]dCTP was added to the PCR reaction together with the standard dNTPs concentration (200 µM each). The PCR reaction product was purified using the QIAquick PCR purification kit (Qiagen) and Chroma Spin TE-200 columns (Clontech). Purified DNA was quantitated by comparing the radioactive DNA fragment with known amounts of a cold PCR product on an agarose gel stained with GelRed (Biotium).

The HindIII digest of λ DNA (New England Biolabs) was labeled at the 3ꞌ end with [α-^32^P]dATP (Perkin Elmer) and the Klenow fragment of DNA polymerase I (New England Biolabs). Free nucleotides were removed with Micro Bio-Spin P-30 Tris chromatography columns (Biorad). Prior to each experiment, the substrate was denatured by heating at 95°C for 5 min to obtain ssDNA. For the preparation of the plasmid-length DNA substrates, the pAttP-S vector, respective annealed oligonucleotides with modified ends and ΦC31 integrase were reacted to obtain the desired DNA fragments (Pinto et al., 2018). Plasmid-length dsDNA substrates with quadruple blocked, 5ꞌ blocked or 3ꞌ blocked DNA ends were generated using the oligonucleotides PC210 and PC211, PC206 and PC207, PC208 and PC209, respectively (Anand et al., 2016). Subsequently, prior to experiments, each substrate was heated at 95°C for 5 min to obtain ssDNA. For the ATPase assay, the 10.3-kbp-long pFB-MBP-hMLH3 plasmid (Ranjha et al., 2014) was linearized with NheI restriction enzyme (New England Biolabs) and the reaction product was purified with QIAquick PCR purification kit (Qiagen). The substrate was denatured at 95°C for 5 min before each experiment.

### Nuclease assays

Nuclease assays with PCR-based DNA substrates were performed in 15 µl volume in 25 mM Tris-acetate pH 7.5, 2 mM magnesium acetate, 1 mM ATP, 1 mM DTT, 0.1 mg/ml BSA (New England Biolabs), 1 mM phosphoenolpyruvate (PEP), 80 U/ml pyruvate kinase (Sigma), 50 mM NaCl and 1 nM substrate (in molecules). Human RPA was included as indicated to saturate all ssDNA. Additional recombinant proteins were then added on ice and the reactions were incubated at 37°C as indicated to perform kinetic experiments. Reactions were stopped by adding 5 µl of 2% stop solution (150 mM EDTA, 2% sodium dodecyl sulfate, 30% glycerol, bromophenol blue) and 1 µl of proteinase K (Sigma) and incubated at 37°C for 10 min. Samples were analyzed by 1% agarose gel electrophoresis. Gels were dried on DE81 chromatography paper (Whatman), exposed to storage phosphor screens (GE Healthcare) and scanned by a Typhoon 9500 phosphorimager (GE Healthcare). Analysis of ssDNA degradation was performed with 0.15 nM (in molecules) 3’-labeled λ DNA/HindIII fragments that were heat-denatured for 5 min at 95°C before adding to the reaction mixture. Differently from above, the reaction buffer contained 3 mM magnesium acetate, no salt, and, unless indicated otherwise, reactions were incubated at 37°C for 8 min. When yeast proteins were used, the experiment was performed at 25°C for 1 min and yeast RPA was added in each reaction. Nuclease assays with ^32^P-labeled Y-shaped DNA substrate (0.1 nM in molecules) were carried out in a similar buffer containing 100 mM NaCl at 37°C and the reactions were stopped by adding 0.5 µl ethylenediaminetetraacetic (0.5 M EDTA) and 1 μl Proteinase K, and incubated at 50°C for 30 min. An equal amount of formamide dye (95% [v/v] formamide, 20 mM EDTA, bromophenol blue) was added and samples were heated at 95°C for 4 min and separated on 15% denaturing polyacrylamide gels (ratio acrylamide:bisacrylamide 19:1, Biorad). After fixing in a solution containing 40% methanol, 10% acetic acid and 5% glycerol for 30 min the gels were dried on 3 mm paper (Whatman) and analyzed as described above. Endonuclease assays with MRN and phosphorylated CtIP (15 µl volume) were performed in nuclease buffer containing 25 mM Tris-HCl pH 7.5, 5 mM magnesium acetate, 1 mM manganese acetate, 1 mM ATP, 1 mM DTT, 0.25 mg/ml BSA, 1 mM phosphoenolpyruvate, 80 U/ml pyruvate kinase, and 1 nM oligonucleotide-based DNA substrate (in molecules). Biotinylated DNA ends were blocked by adding 15 nM streptavidin and incubating the samples 5 min at room temperature. Samples were then processed and analyzed as described above with 15% denaturing gels (Anand et al., 2016; Pinto et al., 2018). Where unlabeled pAttP-S based DNA substrates were used, the reaction buffer was prepared with no salt, but with 3 mM magnesium acetate and 30 nM streptavidin to block biotinylated DNA ends as indicated. Reactions were incubated at 37°C for 30 min and DNA was visualized by staining with GelRed (Biotium).

### Helicase assays

Helicase assays (15 µl volume) were performed in a reaction buffer (25 mM Tris-acetate pH 7.5, 5 mM magnesium acetate, 1 mM ATP, 1 mM DTT, 0.1 mg/ml BSA, 1 mM PEP, 80 U/ml pyruvate kinase and 50 mM NaCl) with 0.1 nM of oligonucleotide-based DNA substrate (in molecules). Recombinant proteins were added as indicated. Unless specified otherwise, reactions were incubated at 37°C for 30 min and stopped as described for nuclease assay with PCR-based DNA substrate. To avoid re-annealing of the substrate, the 2% stop solution was supplemented with a 20-fold excess of the unlabeled oligonucleotide with the same sequence as the ^32^P labeled one. The products were separated by 10% polyacrylamide gel electrophoresis, dried on 17 CHR chromatography paper (Whatman) and analyzed as defined for nuclease assays.

### ATPase assays

The ATPase assays were performed in 25 mM Tris-acetate (pH 7.5), 3 mM magnesium acetate, 1 mM DTT, 0.1 mg/ml BSA, 1 mM ATP, 1 nM of [γ-^32^P] adenosine 5ꞌ-triphosphate (Perkin Elmer) and 0.32 nM (in molecules) of the heat-denatured plasmid-based DNA substrate. RPA and recombinant proteins were added on ice and samples were incubated at 37°C for 10 min. Reactions were stopped with 1.1 µl of 0.5 M EDTA and separated using TLC plates (Merk) and 0.3 M LiCl and 0.3 M formic acid as mobile phase. Dried plates were exposed to storage phosphor screens (GE Healthcare) and scanned by a Typhoon 9500 phosphorimager (GE Healthcare).

### Protein interaction assays

To test for interactions between DNA2 and phosphorylated CtIP, his-DNA2-FLAG was expressed in *Sf*9 cells, cells were lysed and soluble extract containing his-DNA2-FLAG was bound to M2 anti FLAG affinity resin (50 µl, Sigma). The resin was washed with wash buffer 1 (50 mM Tris-HCl pH 7.5, 300 mM NaCl, 10% glycerol, 1 mM PMSF, 0.5 mM β-mercaptoethanol, 0.1% NP40) and incubated for 1 h at 4°C with recombinant purified pCtIP (1 µg) in IP buffer (25 mM Tris-HCl pH 7.5, 0.5 mM DTT, 3 mM EDTA, 100 mM NaCl, 0.20 µg/µl BSA). The resin with bound proteins was washed 5 times with wash buffer 2 (50 mM Tris-HCl pH 7.5, 0.5 mM DTT, 3 mM EDTA, 80 mM NaCl, 0.1% NP40), and proteins were eluted with wash buffer 2 (70 µl) containing 150 ng/µl of FLAG peptide (APExBIO). As a negative control, purified pCtIP was incubated with the resin that had not been bound to his-DNA2-FLAG. The proteins in the eluate were analyzed by western blotting using anti-FLAG primary antibody (Sigma, F3165 diluted 1:1000 in 5% milk in TBS-T) against DNA2-FLAG or anti-CtIP primary antibody (Active Motif, 61141 diluted 1:1000 in blocking solution) using standard procedures.

### Magnetic tweezers assay

The DNA construct for single-molecule experiments containing a 40 nt flap at an adjacent 38 nt gap (see Fig. 4A) was prepared as described before (Kasaciunaite et al., 2019; Levikova et al., 2013; Pinto et al., 2016). To produce the main 6.6 kbp fragment, pNLrep plasmid (Luzzietti et al., 2011) was digested with BamHI and BsrGI restriction enzymes. Simultaneously, a 63 nt gap was created by codigesting five 15 nt spaced BbvCI sites using the nicking enzyme Nt.BbvCI. Into the gap a 65 nt long oligonucleotide possessing a 25 nt complementary sequence and a 40 nt long polithymidine tail was hybridized. In a subsequent ligation step the oligomer was ligated at its 3ꞌ end inside the gap. Also 600 bp long digoxigenin- and biotin-modified handles were ligated with corresponding sticky ends to the termini of the 6.6 kbp fragment.

Single-molecule experiments were carried out in a custom-made magnetic tweezers setup (Huhle et al., 2015; Klaue and Seidel, 2009). The DNA constructs were attached at their biotinylated end to 2.8 µm streptavidin-coated magnetic beads (Dynabeads M280, Thermo Fischer Scientific) and flushed into the fluidic cell, whose bottom slide was covered with antidigoxigenin. After a short incubation to allow the attachment of the digoxigenin-modified DNA end, the excess of the magnetic beads was washed away. Lowering the magnets of the setup allowed to stretch DNA molecules tethered to the magnetic beads. Tracking of the magnetic bead position was performed at 300 Hz using video microscopy and real-time GPU-accelerated image analysis (Huhle et al., 2015). Magnetic forces were calibrated using fluctuation analysis (Daldrop et al., 2015). The DNA unwinding experiments were performed at 37 °C in reaction buffer containing 25 mM Tris-acetate pH 7.5, 5 mM magnesium acetate, 1 mM ATP, 1 mM DTT and 0.1 mg/ml BSA. 2 mM magnesium acetate was used when WRN was present in the reactions. DNA unwinding was initiated by adding equimolar mixtures of 25 nM RPA, 25 nM DNA2 D277A, 25 nM pCtIP and/or 25 nM BLM or WRN (as indicated in the text) in reaction buffer into the fluidic cell. For temperature control of the setup an objective heater (Okolab, Pozzuoli, Italy) was employed. Analysis of unwinding velocities and processivities was carried out using a custom-written MATLAB program (Kemmerich et al., 2016). Errors of obtained rates and processivities are given as standard error of the mean (SEM) throughout.

### SSA reporter assay

0.4 × 10^5^ SA-GFP U2OS cells were seeded in 0.5 ml antibiotic-free media onto 24-well plates with 5 pmol siRNA (siCTRL, 5ꞌ -TGGTTTACATGTCGACTAA; siCtIP, 5ꞌ - GCTAAAACAGGAACGAATC) and 1.8 µl RNAiMAX (Invitrogen) in 100 µl Opti-MEM (Gibco) and cultured for 20 h. The cells were transfected with 0.4 µg I-SceI (pCBASce) and 0.2 µg empty vector (pcDNA3.1) with Lipofectamine 2000 (Invitrogen) in 100 µl Opti-MEM and 0.5 ml antibiotic-free media for 3 hr. After the I-SceI transfection, cells were washed and treated with MRE11 inhibitors 50 µM PFM03 and 50 µM PFM39 (from Davide Moiani and John Tainer, MD Anderson Cancer Center)(Shibata et al., 2014) or mock-treated with DMSO and cultured for 3 days to allow for the repair of the DSB. Cells were then scored for percentage of GFP-positive cells on a LSR Fortessa flow cytometer (BD).

## Supplemental experimental procedures

### Cloning, expression and purification of recombinant proteins

The human EXO1 gene was amplified by PCR using HEXO1FO and HEXO1RE primers to introduce BamHI and XmaI restriction sites as well as C-terminal FLAG tag. The PCR product was digested with BamHI and XmaI restriction endonucleases (New England Biolabs) and ligated into pFastBac1 (Invitrogen), generating pFB-EXO1-FLAG. Human EXO1 was expressed in *Sf*9 insect cells in SFX Insect serum-free medium (Hyclone) using the Bac-to-Bac expression system (Invitrogen), according to manufacturer’s recommendations. Frozen *Sf*9 pellet from 800 ml culture was resuspended in lysis buffer (50 mM Tris-HCl pH 7.5, 0.5 mM β-mercaptoethanol, 1 mM EDTA, 1:400 protease inhibitor cocktail [Sigma], 20 µg/ml leupeptine [Merck Millipore], 0.5 mM PMSF) and incubated at 4°C for 10 min. Glycerol was added to a final concentration of 25%, NaCl was added to a final concentration of 305 mM and the solution was incubated at 4°C for 30 min. The mixture was centrifuged at 55000 g at 4°C for 30 min. The soluble extract was incubated with M2 anti FLAG affinity resin (Sigma) at 4°C for 30 minutes. FLAG resin was washed with TBS wash buffer (20 mM Tris-HCl pH 7.5, 0.5 mM β-mercaptoethanol, 0.5 mM PMSF, 1 mM EDTA, 1:1000 protease inhibitor cocktail, 6.7 µg/ml leupeptine, 10% glycerol, 150 mM NaCl) supplemented with 0.1 % NP40. The last wash was performed with the same buffer but without NP40. Protein was eluted using TBS wash buffer without NP40 supplemented with 200 µg/ml 3XFLAG peptide (Sigma). Peak fractions, as estimated by the Bradford method, were pooled and diluted by adding 1.5 volumes of dilution buffer (50 mM Tris-HCl pH 7.5, 5 mM β-mercaptoethanol, 0.5 mM PMSF, 6.7 µg/ml leupeptine, 10% glycerol). The diluted fractions were loaded on HiTrap SP HP cation exchange chromatography column (GE Healthcare) (0.8 ml/min) and washed with buffer A (50 mM Tris-HCl pH 7.5, 5 mM β-mercaptoethanol, 10% glycerol, 75 mM NaCl). Proteins were eluted using a 5 ml gradient of 75 mM to 1 M NaCl in 0.8 ml fractions. Peak fractions were pooled, snap-frozen in liquid nitrogen and stored at −80°C.

## Acknowledgements

We thank Dr. Davide Moiani and Dr. John Tainer (MD Anderson Cancer Center) for providing MRE11 inhibitors. We thank members of the Cejka laboratory (A. Acharya, G. Reginato, E. Cannavo, A. Sanchez, S. Halder) for critical comments on the manuscript. This work was supported by the European Research Council (ERC, HRMECH, 681630) and the Swiss National Science Foundation (31003A_175444).

## Competing interests

The authors declare no conflict of interest.

---

**Table S1.**
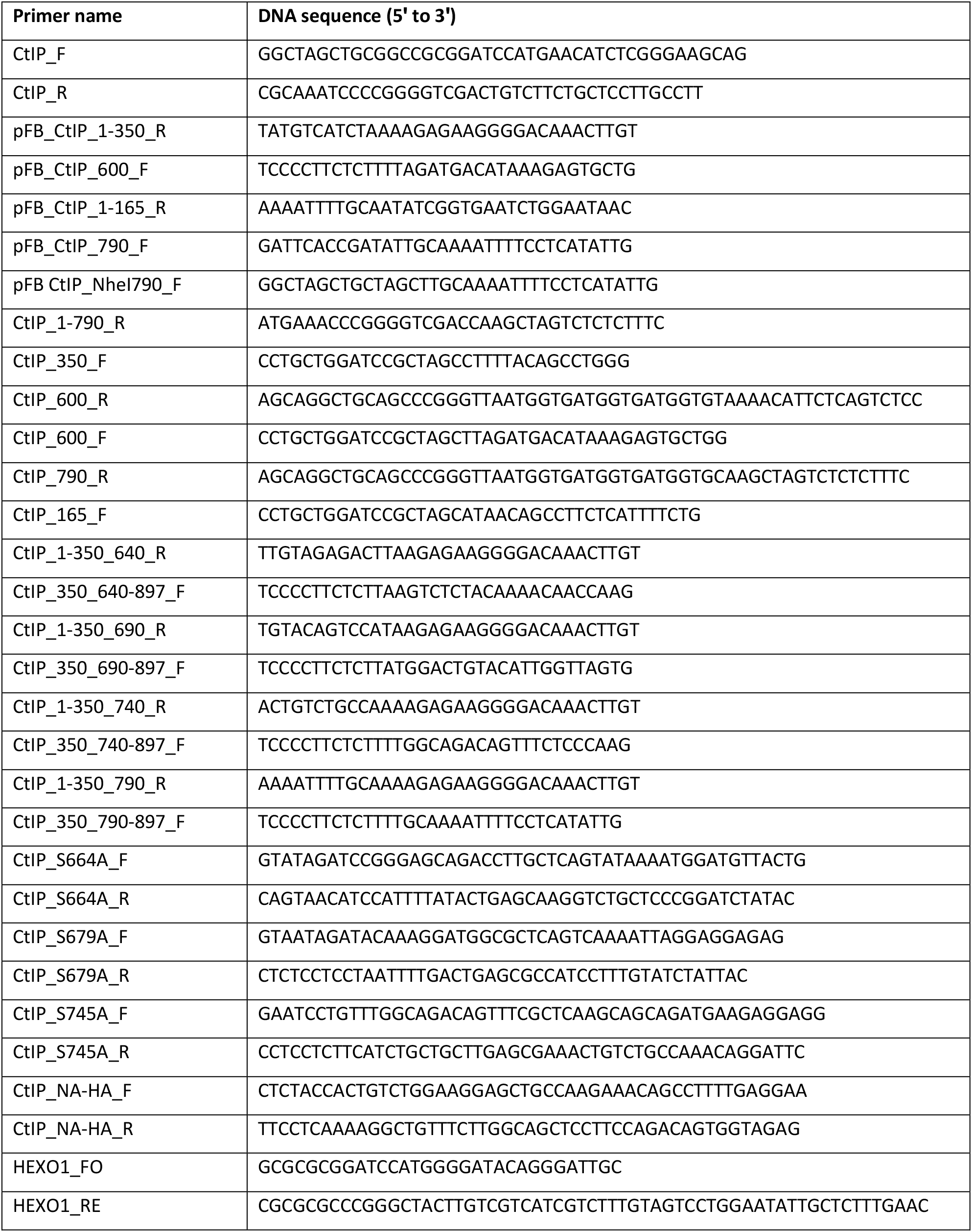
List of oligonucleotides used for cloning and site-directed mutagenesis in this study.

**Table S2.**
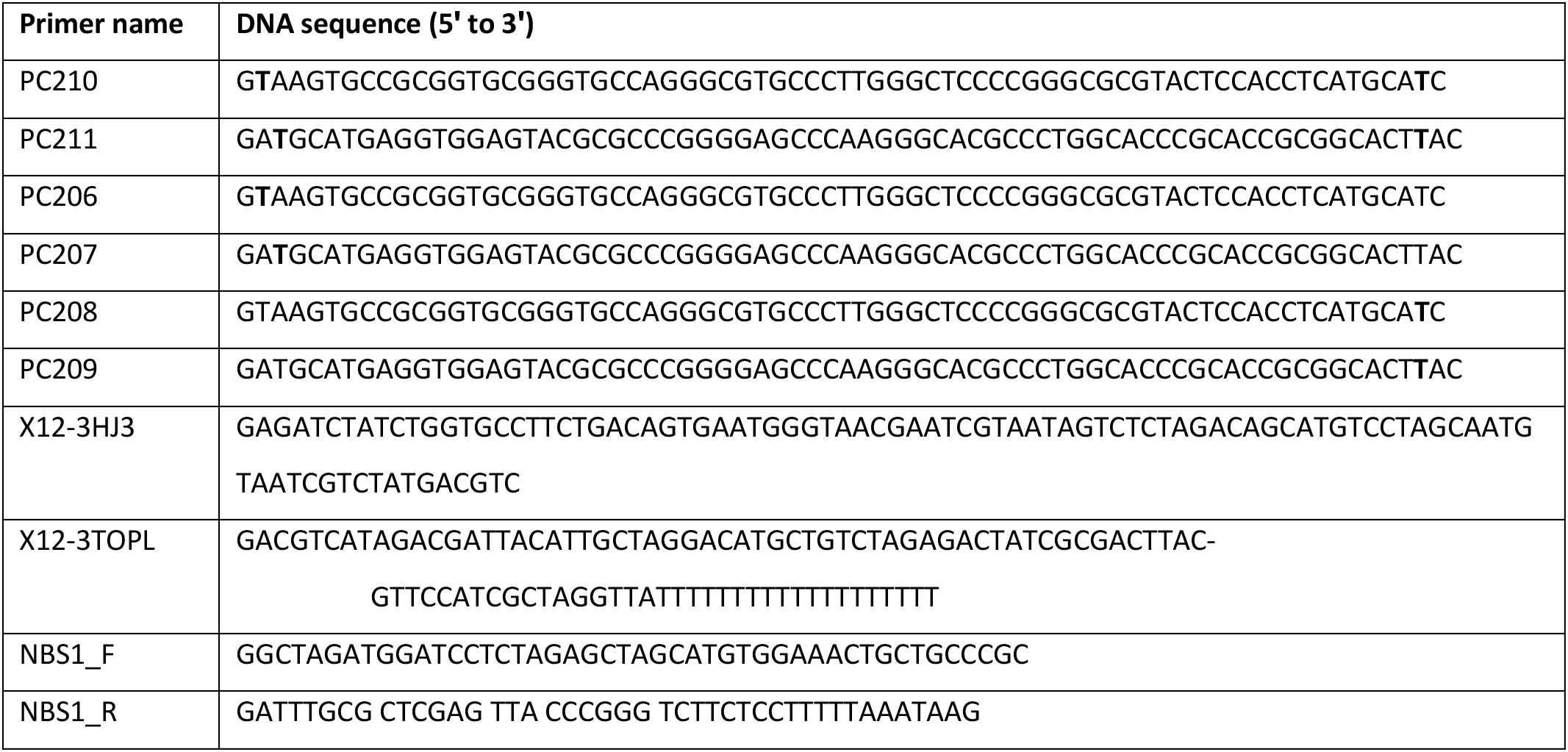
List of oligonucleotides used for oligonucleotide-based DNA substrate in this study. The bold **T** represents the site of the biotin modification.

**Supplementary Figure 1.**
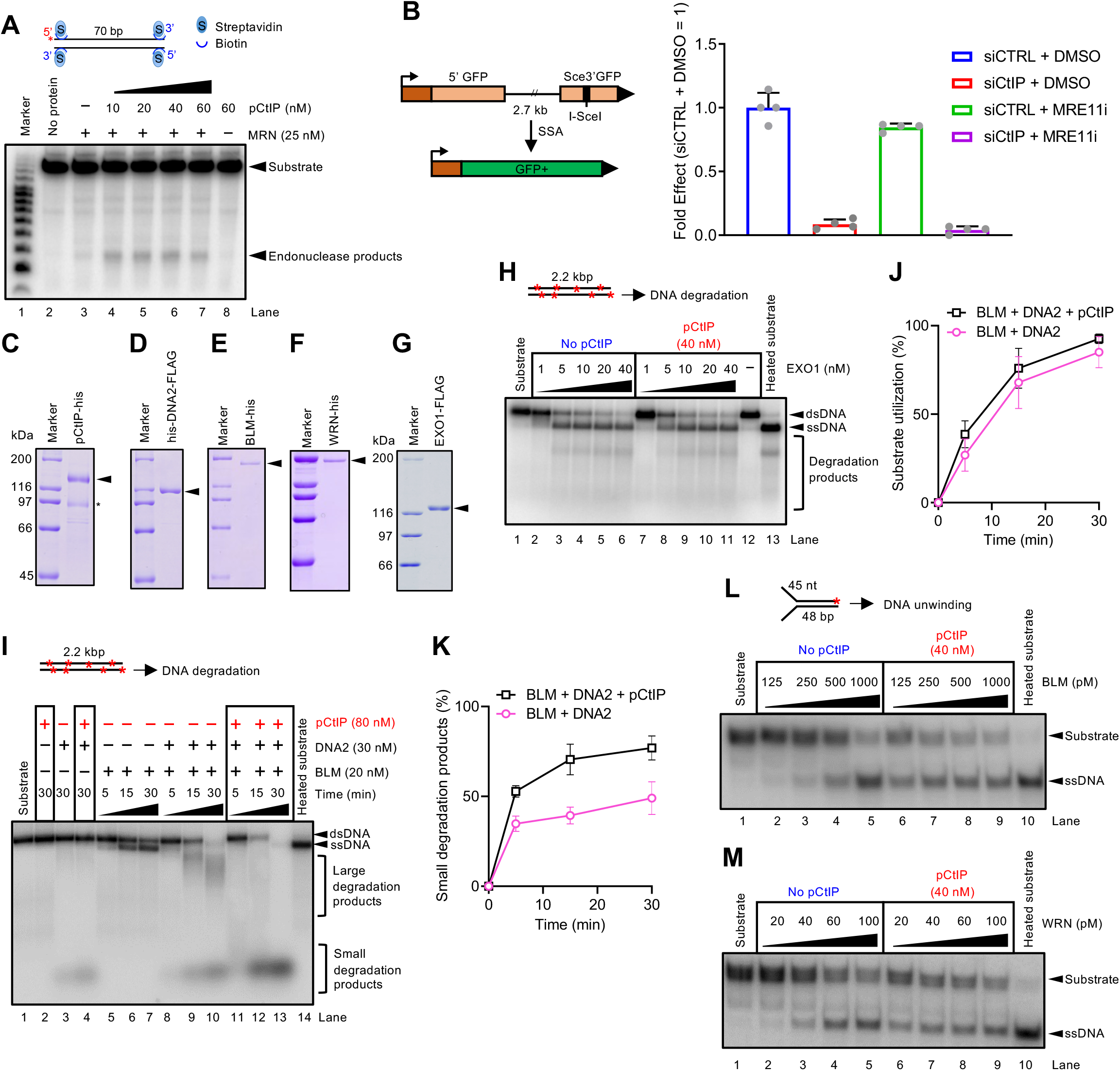
A. Nuclease assay with MRN and various concentrations of phosphorylated CtIP (pCtIP) on a 5’ end-labeled 70 bp-long dsDNA blocked at both ends with streptavidin. Red asterisk indicates the position of the labelling. B. Single-strand annealing reporter assay. CtIP was depleted by siRNA, where indicated, or non-targeting siRNA was used as a negative control (siCTRL). Where indicated, the cells were treated with 50 µM PFM03 and 50 µM PFM39 MRE11 nuclease inhibitors (MRE11i). N = 4; error bars, SD. C-G. Purified proteins used in this study. The polyacrylamide gels were stained with Coomassie Brilliant Blue. * indicates degradation products. H. 2.2 kbp-long dsDNA degradation by EXO1 with or without pCtIP. Panel shows a representative experiment carried out with human RPA (176 nM) and 50 mM NaCl. Red asterisks indicate random labelling. I. DNA end resection by BLM, DNA2 and human RPA (176 nM) in the absence or presence of pCtIP. Panel shows a representative experiment performed with 50 mM NaCl. Red asterisks indicate random labelling. J. Quantitation of overall substrate utilization from experiments such as shown in panel I. N=3; error bars, SEM. K. Quantitation of small degradation products from experiments such as shown in panel I. N=3; error bars, SEM. L. Unwinding of Y-structured DNA by BLM with or without pCtIP, in the presence of human RPA (7.5 nM) and 50 mM NaCl. Moderate DNA stimulation of DNA unwinding by pCtIP was observed. Red asterisk indicates the position of the labelling. M. Experiment as in L, but with WRN. No stimulation of DNA unwinding by pCtIP was observed.

**Supplementary Figure 2.**
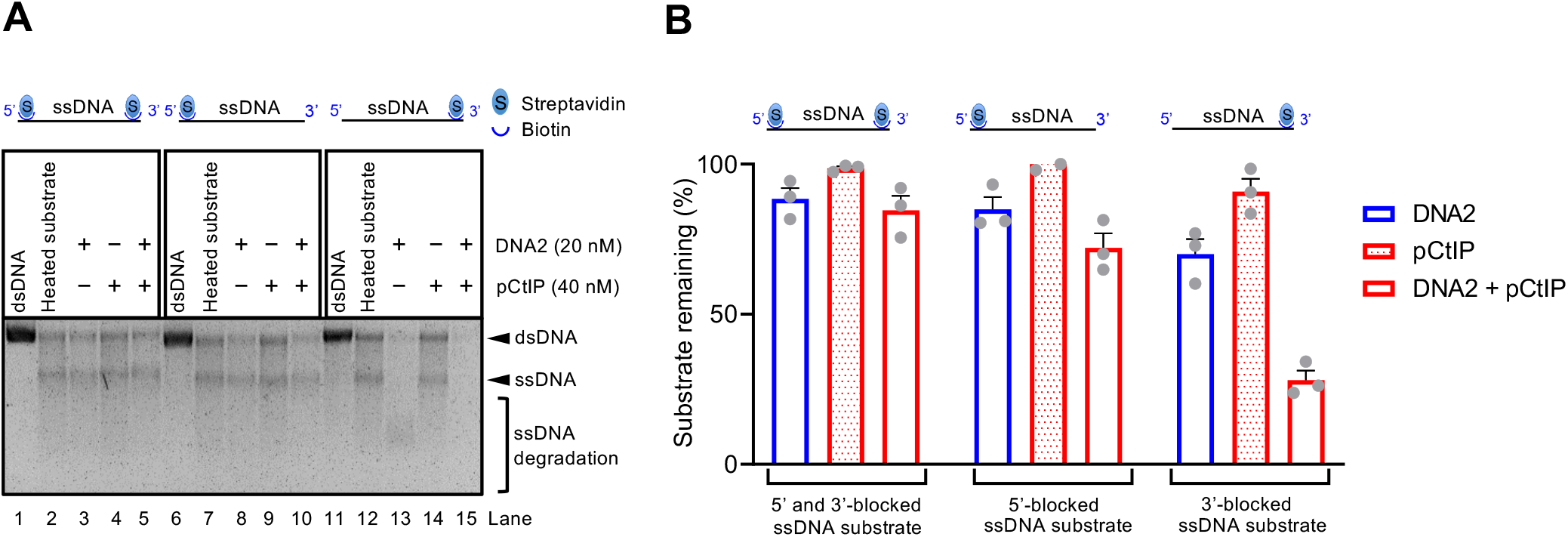
A. 2.7 knt long ssDNA substrates blocked with streptavidin at both ends (lanes 1-5), at the 5’ end (lane 6-10) or at the 3’ end (lanes 11-15), were reacted at 37°C for 30 minutes with DNA2 and/or CtIP in the presence of human RPA (829.8 nM). The gel was stained with GelRed. B. Quantitation of overall remaining substrate from experiments such as shown in panel A. N=3; error bars, SEM. DNA2 preferentially degrades the ssDNA substrate from the 5’ end.

**Supplementary Figure 3.**
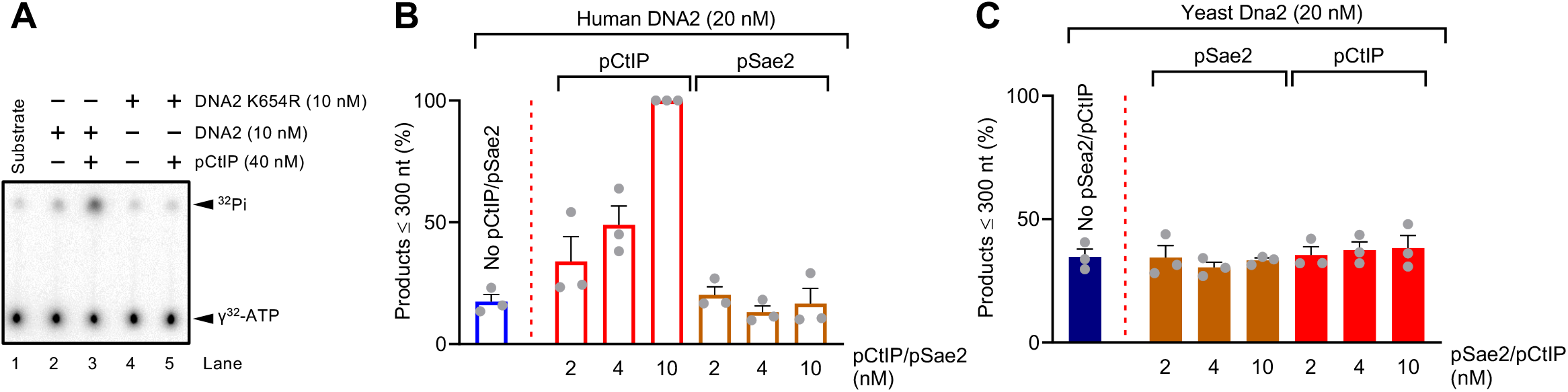
A. ATP hydrolysis by wild type or helicase-deficient K654R DNA2 alone or in the presence of pCtIP. Reactions contained 10.3 kbp-long dsDNA substrate denatured at 95°C for 5 min, and 395.5 nM human RPA. B. Quantitation of small degradation products generated by human DNA2 with pCtIP or pSae2 from experiments such as shown in Fig. 3G. N=3; error bars, SEM. C. Quantitation of small degradation products generated by yeast Dna2 with pSae2 or pCtIP from experiments such as shown in Fig. 3G. N=3; error bars, SEM.

**Supplementary Figure 4.**
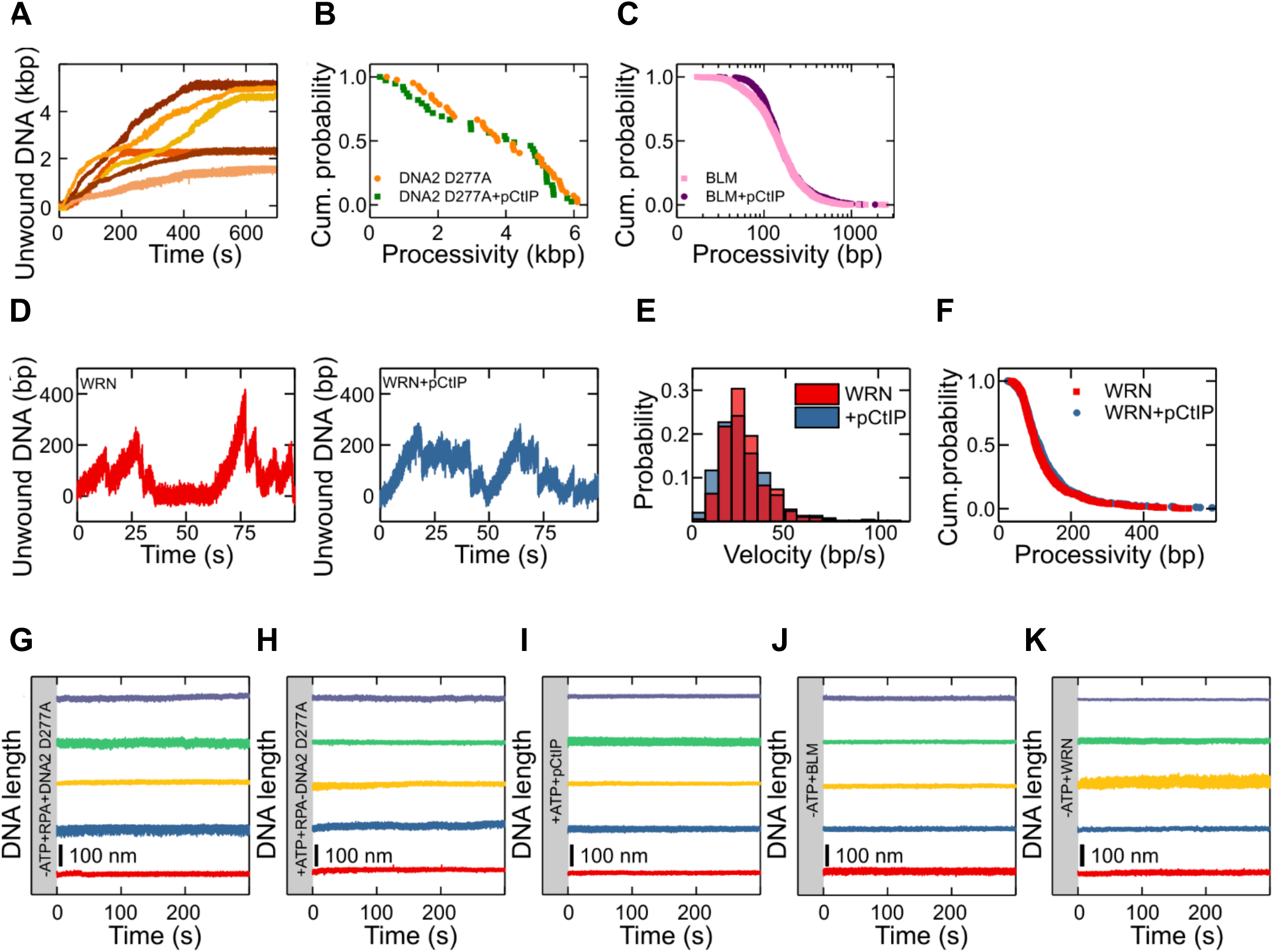
A. Representative DNA unwinding events of nuclease-dead DNA2 (D277A) in the absence of pCtIP as presented in Fig. 4 shown over the full experimental timescale. B. Cumulative probability distributions (shown as survival probability) of the processivity of the individual unwinding events of nuclease dead DNA2 D277A in the absence (orange) and presence (green) of pCtIP with mean values of 3.7±0.3 kbp (SEM) and 3.5±0.3 kbp (SEM), respectively. C. Cumulative probability distributions of the processivity of the individual unwinding events of BLM in the absence (magenta) and presence (purple) of pCtIP with mean values of 179±13 bp (SEM) and 197±15 bp (SEM), respectively. D. Representative DNA unwinding events of WRN in the absence (red) and presence (blue) of pCtIP. E. Histograms of the observed unwinding velocities for WRN in the absence (red) and presence (blue) of CtIP with mean values of 28 ± 2 bp/s (SEM) (N = 751) and 28 ± 3 bp/s (SEM) (N = 547), respectively. F. Cumulative probability distributions of the processivity of the individual unwinding events of WRN in the absence (red) and presence (blue) of pCtIP with mean values of 126 ± 10 bp (SEM) and 134 ± 11 bp (SEM), respectively. G–K. DNA unwinding studied by: (G) DNA2 D277A and human RPA in the absence of ATP, (H) human RPA and ATP in the absence of DNA2 D277A, (I) pCtIP and ATP in the absence of BLM and DNA2 D277A as well as (J) BLM or (K) WRN in the absence of ATP.

**Supplementary Figure 5.**
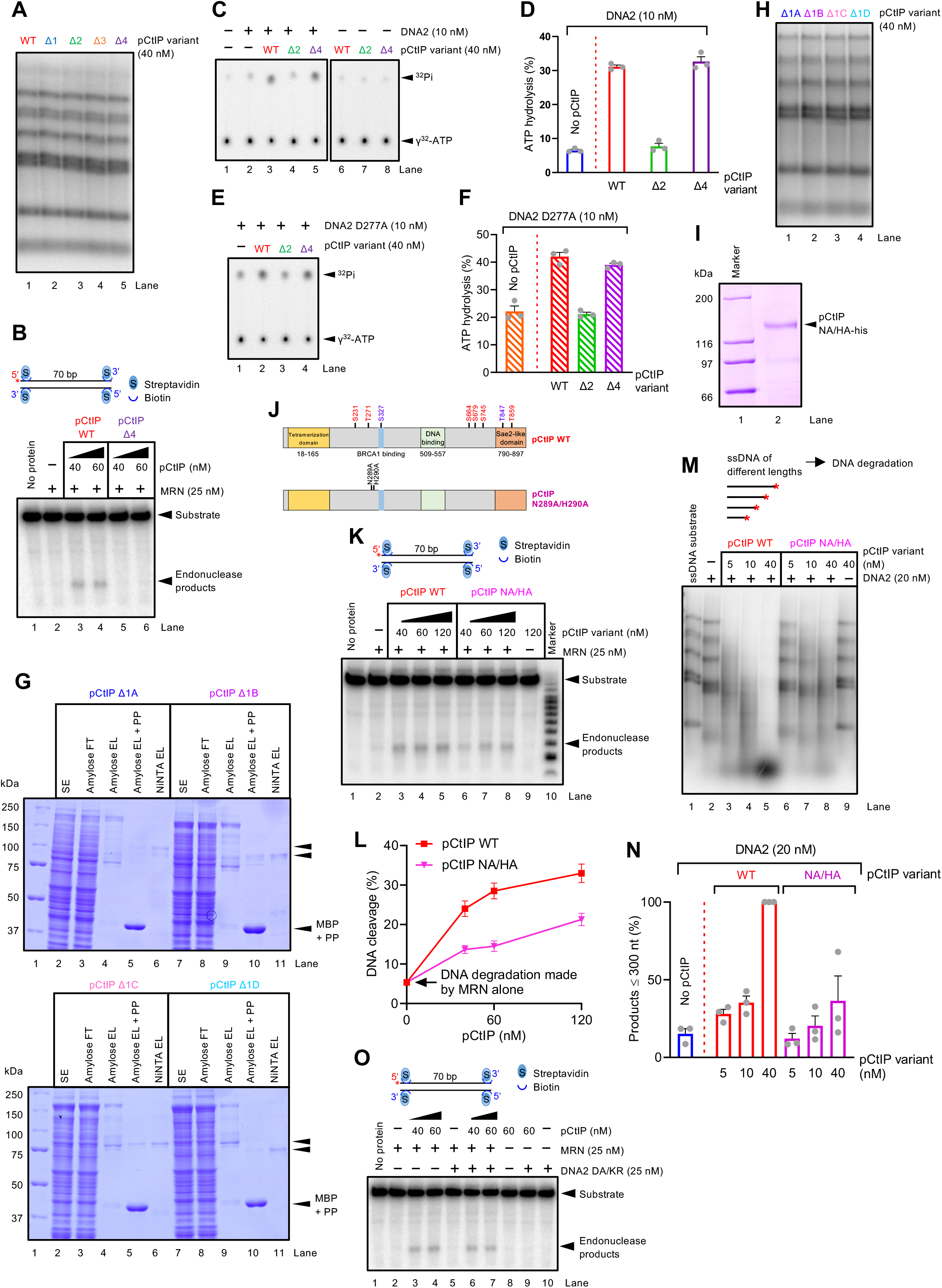
A. Experiment as in Fig. 5B, but carried out only with wild type phosphorylated CtIP or internal deletion variants, in the presence of human RPA (864 nM). B. Endonuclease assay with MRN (25 nM) and wild type full-length or pCtIP Δ4 variants. Red asterisk indicates the position of the labelling. C. ATP hydrolysis by DNA2 without or with pCtIP and its variants. Reactions contained 10.3 knt-long ssDNA, 395.5 nM human RPA and no added salt. The ATPase activity of DNA2 alone or with wild type pCtIP is the same as shown in Fig. 3E and is shown again as a reference. D. Quantitation of data such as shown in panel C. N=3; error bars, SEM. The ATPase activity of DNA2 alone and with wild type pCtIP is the same as in Fig. 3E. E. ATP hydrolysis by nuclease-deficient DNA2 D277A without or with pCtIP variants, as indicated. Reactions contained 10.3 knt-long ssDNA, 395.5 nM human RPA and no salt. F. Quantitation of data such as shown in panel E. N=3; error bars, SEM. G. Gradient (4-15%) polyacrylamide gels (Biorad) stained with Coomassie blue showing fractions from representative purifications of pCtIP Δ1A– Δ1D variants. SE, soluble extract; FT, flow through; EL, eluate. H. Experiment as in Fig. 5G, but carried out only with wild type phosphorylated CtIP or internal deletion variants, in the presence of human RPA (864 nM). I. Recombinant pCtIP NA/HA mutant analyzed on a polyacrylamide gel stained with Coomassie Brilliant Blue. J. A schematic representation of the positions of the mutations in the pCtIP NA/HA variant. pCtIP wild-type is again shown as a reference. K. Endonuclease assay with MRN (25 nM) and various concentrations of wild type pCtIP or the NA/HA mutant. Red asterisk indicates the position of the labelling. L. Quantitation of data such as shown in panel J. N=3; error bars, SEM. M. Degradation of ssDNA fragments of various lengths by DNA2 without or with various concentrations of wild type pCtIP or the NA/HA mutant, in the presence of human RPA (864 nM). Red asterisk indicates the position of the labelling. N. Quantitation of data such as shown in panel L. N=3, error bars, SEM. O. Endonuclease assay with MRN (25 nM) and various concentrations of pCtIP, with or without nuclease-helicase-deficient DNA2 D277A K654R. Red asterisk indicates the position of the labelling.

**Supplementary Figure 6.**
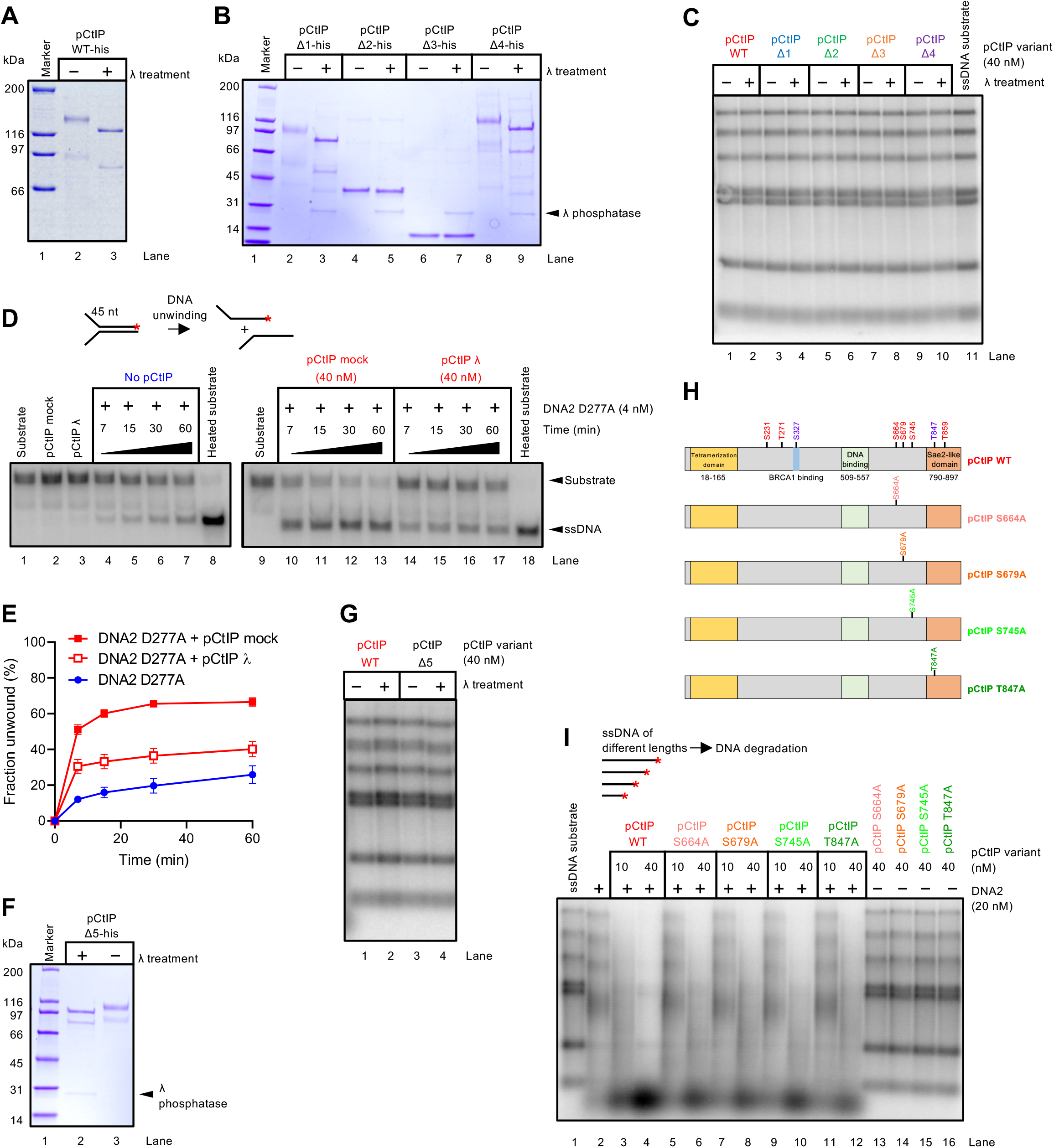
A. Electrophoretic mobility of pCtIP either not-treated (lane 2) or treated (lane 3) with λ phosphatase. The pCtIP variants were separated on a gradient (4-15%) polyacrylamide gel (Biorad) and stained with Coomassie Brilliant Blue. B. Electrophoretic mobility of pCtIP variants Δ1, Δ2, Δ3 or Δ4 either not-treated or treated with λ phosphatase. The CtIP variants were separated on a gradient (4-15%) polyacrylamide gel (Biorad) and stained with Coomassie Brilliant Blue. C. Experiment as in Fig. 6A, but carried out without DNA2. D. A representative experiment showing the unwinding activity of nuclease-deficient DNA2 D277A without or with mock-treated or λ-treated pCtIP on an oligonucleotide-based Y-structured (45 nt/48 bp) DNA substrate. Reactions were supplemented with human RPA (7.5 nM) and 50 mM NaCl, and analyzed by electrophoresis in a native 10% polyacrylamide gel. Red asterisk indicates the position of the labelling. E. Quantitation of data such as shown in panel D. N=3, error bars, SEM. F. Electrophoretic mobility of pCtIP variant Δ5 either treated (lane 2) or not-treated (lane 3) with λ phosphatase. G. Experiment as in Fig. 6D, but carried out without DNA2. H. A schematic representation of purified recombinant non-phosphorylatable pCtIP variants (S664A, S679A, S745A, T847A). pCtIP wild type is again shown as a reference. I. Degradation of ssDNA fragments of various length with DNA2 alone (20 nM) or with pCtIP wild type or non-phosphorylatable variants, in the presence of human RPA (864 nM). Red asterisk indicates the position of the labelling.

